# FtsZ assembles the bacterial cell division machinery by a diffusion-and-capture mechanism

**DOI:** 10.1101/485656

**Authors:** Natalia Baranova, Philipp Radler, Víctor M. Hernández-Rocamora, Carlos Alfonso, Mar López-Pelegrín, German Rivas, Waldemar Vollmer, Martin Loose

## Abstract

The mechanism of bacterial cell division is largely unknown. The protein machinery performing cell division is organized by FtsZ, a tubulin-homolog that forms treadmilling filaments at the cell division site. Treadmilling is thought to actively move proteins around the cell thereby distributing peptidoglycan synthesis to make two new cell poles. To understand this process, we reconstituted part of the bacterial cell division machinery using the purified components FtsZ, FtsA and truncated transmembrane proteins essential for cell division. We found that membrane-bound cytosolic peptides of FtsN and FtsQ co-migrated with treadmilling FtsZ-FtsA filaments. Remarkably, rather than moving in a directed fashion, individual peptides followed FtsZ filaments by a diffusion-and-capture mechanism. Our work provides a mechanism for how the Z-ring dynamically recruits divisome proteins and highlights the importance of transient interactions for the self-organization of complex biological structures. We propose that this mechanism is used more widely to organize and transmit spatiotemporal information in living cells.

**One Sentence Summary:** FtsZ treadmilling assembles bacterial division machinery by diffusion-and-capture mechanism.

## Introduction

Cells propagate by cytokinesis. Most bacteria accomplish cell fission by assembling a dynamic protein complex, called the divisome, which spans from the cytoplasm through the membrane in the plane of division. Divisome assembly is initiated by the polymerization of the tubulin-related GTPase FtsZ into a discontinuous ring-like structure (the Z-ring) at the nascent division site. The Z-ring is able to recruit about 20 additional proteins including several peptidoglycan (PG) synthases and hydrolases. Together, they build the inwards growing cell wall and concomitantly separate the daughter cells^1^. It has been observed that FtsZ forms treadmilling filaments on supported bilayers *in vitro*^2^ and in live cells where they circle around the cell division site^3,4^. In a treadmilling filament, monomers are added to one end of the filament and detach from the opposite end, resulting in the apparent forward movement of the polymer while the individual monomers remain static. Importantly, the directed motion of the transmembrane PG synthases PBP3 (FtsI) in *E. coli*^4^ and PBP2b in *B. subtilis*^3^ was found to correlate with FtsZ treadmilling, suggesting that FtsZ filaments actively move these transmembrane proteins of the divisome and provide spatiotemporal control over the generation of new cell poles. However, these studies did not provide a mechanistic model for how stationary subunits within the treadmilling FtsZ protofilament can facilitate the directed motion of PG synthases^5^. Therefore, the molecular processes allowing the Z-ring to organize the components of the divisome in space and time remained unknown.

FtsZ filaments are attached to the inner leaflet of the cytoplasmic membrane via the membrane anchors ZipA and FtsA. Of those two proteins, the latter appears to be most important as ZipA can be bypassed^6,7^. FtsA appears to be essential in *E. coli*, indicating that it may also be required for a signaling role triggering later steps in cytokinesis^8,9^. In fact, FtsA was previously suggested to recruit other essential cell division proteins including FtsL, FtsN, FtsQ and PBP3 to the Z-ring (Fig. 1a, left)^7,10–13^. However, most of these data were obtained *in vivo* and could not reveal whether or not the protein-protein interactions were direct.

**Figure 1.**
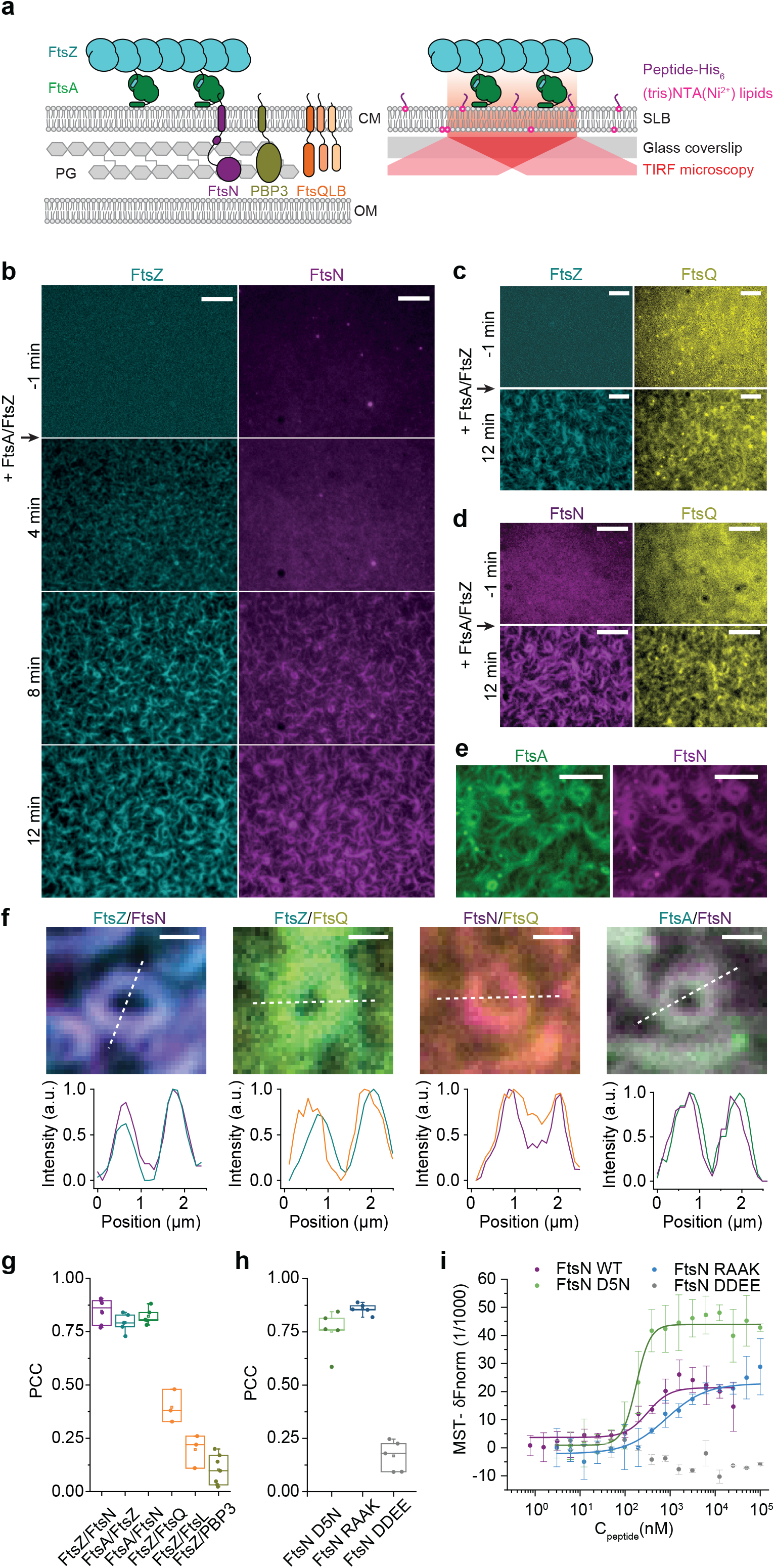
FtsZ/FtsA filaments co-localize with cytoplasmic tails of FtsN and FtsQ. a. Left side: Illustration of key cytosolic interactions and selected divisome proteins with cytosolic tails, FtsN, PBP3 and FtsQLB. Signaling from FtsZ/FtsA co-filaments to the peptidoglycan (PG) layer between the cytoplasmic (CM) and outer membrane (OM). FtsA interacts with the N-terminal cytoplasmic peptides of the transmembrane proteins FtsN, PBP3 and the protein complex FtsQLB. Right side: Illustration of the *in vitro* reconstitution approach used to study interactions between the cytoskeleton and downstream cell division proteins. The N-terminal peptides are attached to the surface of a supported lipid bilayer (SLB) on a glass coverslip.
b. Starting from a homogeneous distribution, CF488-FtsN_cyto_His (magenta) is organized into filaments by the addition of FtsA and Cy5-FtsZ (cyan) at t = 0 min. Scale bars are 5 μm. Supplementary Video 1.
c. Cy5-FtsQ_cyto_His (yellow) is sorted into filaments by the addition of FtsA and Alexa488-FtsZ (cyan) at t = 0 min. Scale bars are 5 μm. Supplementary Video 2.
d. Two peptides Alexa488-FtsN_cyto_His (magenta) and Cy5-FtsQ_cyto_His (yellow) simultaneously present on the membrane (-1 min) are organized into similar dynamic pattern after the addition of unlabeled FtsA and FtsZ proteins (12 min). Scale bars are 5 μm. Supplementary Video 3
e. Dual-color TIRF imaging FtsN_cyto_His is organized into a cytoskeleton pattern via direct binding to FtsA. Micrographs of the membrane-bound Cy5-FtsN_cyto_His (magenta) colocalized with TMR-FtsA (green). Scale bars are 5 μm. Supplementary Video 4.
f. Top row: Enlarged micrographs of ring-like structures formed by co-assembled proteins: Cy5-FtsZ/CF488-FtsN_cyto_His, Alexa 488-FtsZ/Cy5-FtsQ_cyto_His, Alexa488-FtsN_cyto_His/Cy5-FtsQ_cyto_His and TMR-FtsA/Cy5-FtsN_cyto_His. Bottom row: Corresponding intensity profiles across the diameter of the ring taken along the dashed lines. Scale bars are 1 μm.
g. Colocalization of membrane-bound peptides as quantified by the Pearson Correlation coefficient (PCC). FtsN_cyto_His strongly colocalizes with FtsZ (PCC = 0.86 ± 0.05; n = 5, magenta) and FtsA (PCC = 0.80 ± 0.03; n = 5, green), comparable to FtsA/FtsZ colocalization (0.79 ± 0.04; n = 5, cyan). The colocalization of FtsZ with FtsQ_cyto_His is weaker (0.40 ± 0.06; n = 3, light orange, p-value = 6 × 10^−5^) and further decreased for FtsL_cyto_His-FtsZ (0.20 ± 0.06; n = 3, dark orange, p-value = 8 × 10^−6^) and PBP3_cyto_His-FtsZ (0.09 ± 0.07; n = 7, dark yellow, p-value = 6 × 10^−10^). Each dot represents an independent experiment.
h. Colocalization of FtsN_cyto_His with FtsZ/FtsA cofilaments is specific. Colocalization efficiency with FtsZ is comparable between FtsN_cyto_His, FtsN_cyto-D5N_His (PCC = 0.75 ± 0.09; light green, n = 5, p = 0.07) and FtsN_cyto-RAAK_His (PCC = 0.86 ± 0.02; n = 5, blue, p = 0.97). In contrast, for FtsN_cyto-_DDEEHis (PCC =0.17 ± 0.06; n = 5, gray, p-value = 2 × 10^−7^) it is significantly lower.
i. FtsN peptides bind to FtsA with different affinities. Interaction of FtsN peptides with FtsA in solution was measured by Microscale Thermophoresis. FtsN_cyto-D5_NHis (app. K_D_ = 0.16 ± 0.03 μM, n = 3) has a binding affinity comparable to the wild-type FtsN peptide (app. K_D_ = 0.33 ± 0.1 μM, n = 3), while FtsN_cyto-RAAK_His (app. K_D_ = 0.82 ± 0.1 μM, n = 3) shows intermediate affinity and FtsN_cyto-D_DEEHis shows no interaction. Titration curves were fitted with a Hill equation, where the Hill coefficient was set to 1 (solid lines).

FtsL, FtsN, FtsQ, and PBP3 are generally considered to be late recruited proteins, while recent evidence showed that FtsN can also localize early to the division site^14^. All these proteins contain a short N-terminal cytoplasmic tail, a single transmembrane-helix and a larger periplasmic domain. FtsQ, FtsL and FtsB form a complex in the inner membrane which, like FtsN, is thought to activate the PG synthase PBP3^15^. As FtsA binds peripherally to the cytoplasmic membrane, it would be ideally oriented to interact with the cytoplasmic tails of cell division proteins linking cell wall synthesis to Z-ring dynamics^16^. However, important questions remain unanswered: which of these cytoplasmic tails interact with FtsA, how are these interactions affected by FtsA’s role as membrane anchor of FtsZ, and how can cytosolic interactions facilitate the co-migration of cell division proteins with treadmilling FtsZ?

To address these questions, we developed an *in vitro* assay able to recapitulate early stages of cytokinesis dynamics in *Escherichia coli*. We found that the cytoplasmic tails of FtsN and FtsQ, but not PBP3 and FtsL, are recruited to membrane-anchored, treadmilling filaments of FtsZ and FtsA by a diffusion-and-capture mechanism. In this transient assembly, FtsZ, FtsA and cytoplasmic peptides remain stationary, and their accumulations disperse as soon as FtsZ filaments depolymerize. As the treadmilling filament moves forward, the associated proteins follow its effective translational motion giving rise to their apparent movement. Our results support a model in which treadmilling FtsZ filaments initiate and guide cell wall synthesis by continuously increasing the local concentration of key division proteins above a critical threshold needed for septation.

## Results

### The cytoplasmic peptides of FtsN and FtsQ co-localize with FtsZ/FtsA co-filaments on supported membranes

Cell division proteins are distributed homogeneously in the cell membrane in the absence of a Z-ring, but get recruited to the cell division site when FtsZ polymerizes on the membrane^17^. To recapitulate this situation, we used supported lipid bilayers and attached to them the fluorescently labeled cytoplasmic peptides of our candidate divisome proteins FtsL, FtsN, FtsQ and PBP3 (Fig. 1a, right). In the absence of other proteins (Fig1b-d, −1min), the membrane-bound peptides from FtsN, FtsQ, FtsL or PBP3 were homogeneously distributed and diffused freely on these membranes. When we allowed FtsZ and FtsA to polymerize in the presence of a membrane-bound candidate peptide, the fluorescent signal for FtsN_cyto_His and FtsQ_cyto_His increasingly overlapped with the cytoskeletal structures formed by FtsA and FtsZ (Fig. 1b-c, Supplementary Videos 1 and 2), while FtsL_cyto_His and PBP3_cyto_His continued to exhibit a homogeneous membrane-distribution in the presence of FtsZ-FtsA cofilaments (Supplementary Fig. 1a). Remarkably, FtsQ_cyto_His and FtsN_cyto_His formed filament-like patterns even when both peptides were simultaneously present (Fig. 1d, Supplementary Video 3). In all cases, overlap for FtsQ_cyto_His was lower than for FtsN_cyto_His as quantified by the Pearson Correlation Coefficient (PCC) (Fig. 1f, g). Interestingly, while there was a strong colocalization between FtsA and the monomeric FtsNcytoHis on the membrane (Fig. 1e, Supplementary Video 4), previous observations found that only a dimerized form of FtsN_cyto_ interacted with FtsA in solution^18^. To test this apparent discrepancy, we studied the interactions between FtsA and full-length FtsN or the cytoplasmic peptides of divisome proteins either by analytical ultracentrifugation or microscale thermophoresis (MST). While FtsN was able to form oligomers in solution (Supplementary Fig. 1b), full length FtsN and FtsN_cyto_His interacted with FtsA with similar affinity (K_D_ values of 250 ± 51 and 324 ± 96 nM, respectively, Supplementary Fig. 1c-d, Supplementary Table 2). Moreover, FtsQ_cyto_His, FtsL_cyto_His and PBP3_cyto_His showed no interaction with FtsA in solution (Supplementary Fig. S1d). As FtsQ_cyto_His did not bind to FtsA in MST experiments, and only colocalized with FtsZ/FtsA cofilaments when attached to the membrane (Fig. 1 and Supplementary Fig.1 e), we believe that confining the peptide to the supported bilayer could enhance its apparent affinity to FtsA as it is optimally aligned on the membrane surface.

Furthermore, our experiments suggested that ionic interactions play an important role as the interaction between FtsA and FtsN was abolished at increased salt concentrations (300 mM compared to 100 mM KCl) (Supplementary Fig. 1f). To test the ionic nature of the FtsN-FtsA interaction, we studied known mutants of the cytoplasmic peptide of FtsN, where the conserved positively charged residues at positions 16-19 (RRKK) were replaced with more negatively charged (DDEE) or hydrophobic (RAAK) residues^14,18^. The third mutant had an asparagine residue instead of aspartic acid at position 5 (D5N)^7^. FtsN_cyto-D5N_His and FtsN_cyto-RAAK_His showed similar colocalization with FtsZ compared to FtsN_cyto_His (Supplementary Fig. 2a,b) and both mutant peptides interacted with FtsA in solution (Fig. 1i). In contrast, FtsN_cyto-DDE_EHis showed much weaker overlap with FtsZ on the membrane and no interaction in solution (Fig. 1h-i, Supplementary Fig. 2a,b and Supplementary Table 2). To conclude, our assay allows testing interactions between the components of the bacterial cell division machinery, providing evidence for a direct interaction between FtsA and the cytoplasmic tails of FtsN and FtsQ.

### The cytoplasmic tails of FtsN and FtsQ comigrate directionally with treadmilling FtsZ/FtsA co-filaments

Next, we acquired dual-color time-lapse movies to simultaneously image the dynamics of FtsZ and membrane-bound cytoplasmic peptides. We found that both FtsN_cyto_His and FtsQ_cyto_His closely followed the dynamic behavior of FtsZ (Fig. 2a-b). To better visualize the dynamics of the proteins, we constructed differential time-lapse movies, which show the intensity difference between two frames separated by a constant time delay (Fig 2a, Supplementary Fig. 3). This gives rise to fluorescent clusters that represent the amount of protein added to a co-polymer in a given time step, i.e. it specifically visualizes the growth of an FtsZ filament bundle as well as the effective motion of peptide accumulations (Supplementary Fig. 3a-c, Supplementary Video 5). Kymographs of either FtsZ and FtsN_cyto_His or FtsZ and FtsQ_cyto_His along the circumference of rotating rings showed that in both cases the proteins migrated together (Fig. 2a,b and Supplementary Fig. 3d). We continued to characterize the interaction with FtsN_cyto_His because its colocalization with FtsA was much stronger than that of FtsQ_cyto_His. The directed motion of the membrane-bound peptides was abolished when we replaced GTP with the slowly hydrolyzable analog GMPCPP, which does not allow for FtsZ treadmilling and filament reorganization (Fig. 2c & Supplementary Fig. 3e, Supplementary Video 6). Interestingly, in this case the colocalization of FtsN_cyto_His and FtsZ was slightly but significantly decreased, suggesting that FtsZ/FtsA filaments dynamics facilitates the interaction with FtsN_cyto_His (Fig. 2d).

**Figure 2.**
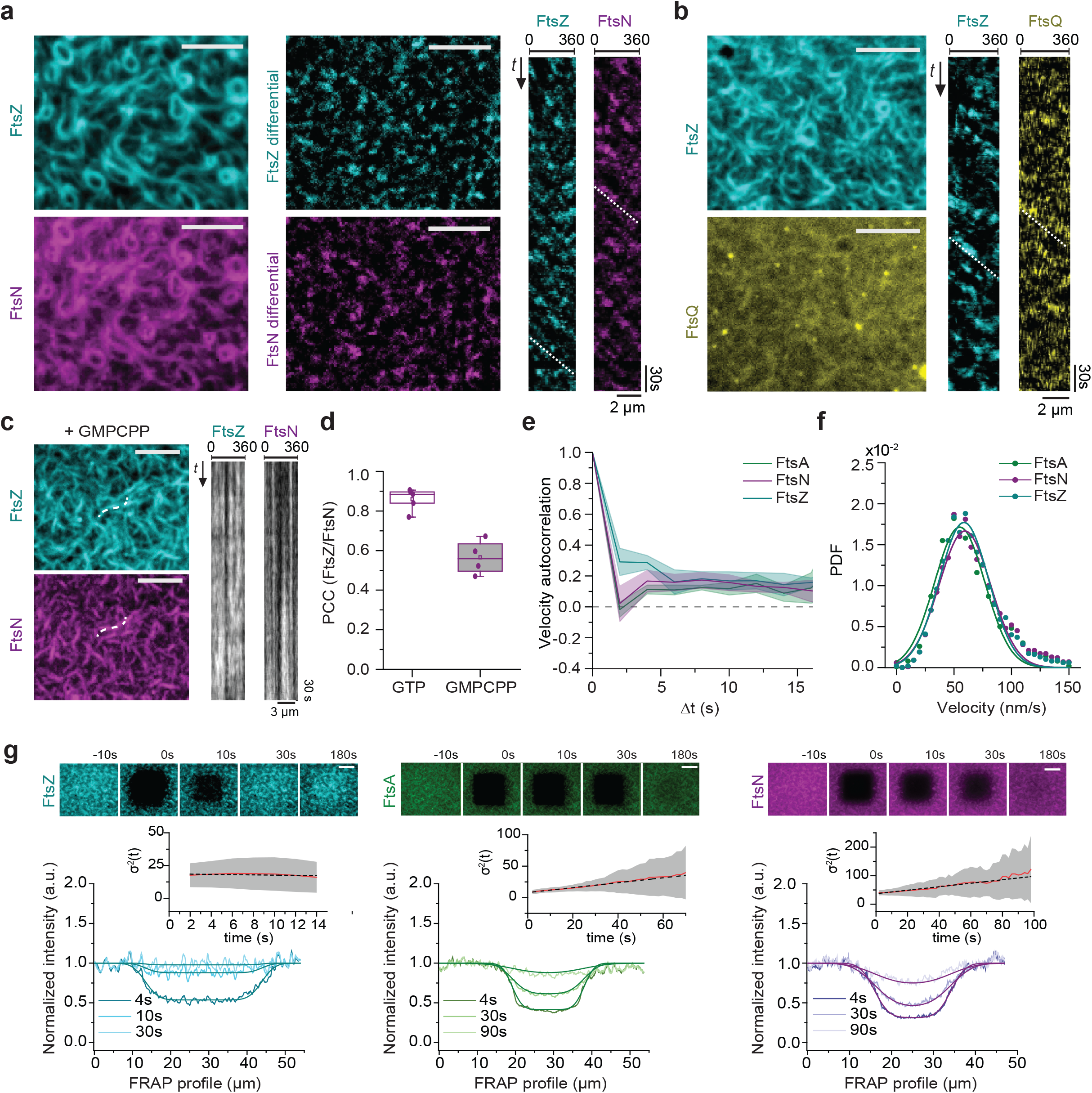
Cytoplasmic tails of FtsN and FtsQ co-migrate directionally with treadmilling FtsA/FtsZ cofilaments and arise from different underlying dynamics. a. FtsNcytoHis follows the dynamic behavior of the FtsZ filaments. Left: Micrographs of Cy5-FtsZ (cyan) and CF488-FtsN_cyto_His (magenta) mirroring FtsZ filaments. Middle: Micrographs showing differential images from the corresponding micrographs. Right: Representative differential kymographs of treadmilling dynamics taken along the contour of a rotating FtsZ ring. Kymographs taken from differential images improve the visualization of FtsZ (v = 60.5 ± 6.0 nm/s; cyan, n = 20) and FtsN_cyto_His (v = 55.5 ± 6.5 nm/s; magenta, n = 20) dynamics. Scale bars are 5 μm.
b. FtsQ_cyto_His follows the dynamic behavior of the FtsZ filaments. Left: Corresponding micrographs of Alexa488-FtsZ (cyan) and Cy5-FtsQ_cyto_His (yellow). Right: Representative kymographs of treadmilling dynamics built along the contour of rotating FtsZ were prepared from the corresponding differential images. FtsQ_cyto_His is moving with the same velocity as FtsZ (v = 54.5 ± 5.9 nm/s; yellow, n = 10). Scale bars are 5 μm.
c. FtsZ treadmilling and FtsN_cyto_His co-migration is GTP-dependent while colocalization is not. TIRF micrographs are revealing colocalized Cy5-FtsZ and CF488-FtsN patterns with GMPCPP (left) and static kymographs (right). Scale bars are 5 μm. Scale bars are 5 μm. Supplementary Video 6
d. Dynamic treadmilling increases the FtsZ/FtsN_cyto_His colocalization. In the presence of GMPCPP the colocalization efficiency is 1.5-fold lower comparing to the presence of GTP (n = 4, p-value = 2.24 × 10^−4^).
e. On the ensemble level all proteins move directionally, as revealed by velocity autocorrelation analysis. Positive values confirm directional motion for FtsZ (cyan), FtsA (green) and FtsN_cyto_His (magenta). Shaded areas represent the standard deviation (n(FtsN) = 8, n(FtsA) = 3; n(FtsZ) = 12).
f. Velocity histograms fitted to Gaussian distributions (solid line) did not reveal any significant difference in the velocity of directed motion of FtsZ (v = 58.7 ± 21.5 nm/s; n = 12; cyan), FtsA (v = 55.0 ± 21.9 nm/s; n = 3; green, p-value = 0.21) and FtsN_cyto_His (v = 59.4 ± 22.4 nm/s; n = 8, magenta, p-value = 0.94). The velocity values represent mean and SD from the corresponding Gaussian fits (R^2^ = 0.94-0.97). PDF stands for Probability density function.
g. Micrographs of FRAP experiments (top) and corresponding intensity profiles across photobleached regions of Cy5-FtsZ (left, cyan), TMR-FtsA (middle, green) and CF488-FtsN_cyto_His (right, magenta) at different time points. The first micrograph shows the area before bleaching, micrographs at 0 s correspond to the pattern at the first frame after bleaching. Scale bars are 10 μm. Fitting a Gaussian error function to the profiles to obtain *σ*^2^(t) points reveals changes in the profile shape (inset plots). For FtsZ, *σ*^2^ remains constant, showing that fast recovery of the photobleached region is dominated by a homogenous exchange of FtsZ monomers. For FtsA and FtsN_cyto_His, *σ*^2^ increases with time, consistent with a strong contribution of lateral diffusion for recovery. This can also be seen by the delayed recovery of fluorescence in the center of the bleached area compared to its edges. See Supplementary Video 7.

Differential movies also allowed us to use particle-tracking methods to analyze migration trajectories of the fluorescent clusters in greater detail. Velocity autocorrelation of treadmilling trajectories for FtsZ, FtsA and FtsN_cyto_His showed positive values for longer time delays, confirming directional motion for all three proteins (Fig. 2e). The rapid drop at short time delays is expected for highly curved trajectories. As motion was directional, we could use the distribution of displacements to calculate the average velocities of FtsZ treadmilling as well as the velocity of co-migration for FtsA and FtsN_cyto_His. We found that all three proteins moved at the same velocity (Fig. 2f and Supplementary Fig. 3f,g). Together, these data show that FtsZ treadmilling driven by GTP hydrolysis is able to power the co-migration of membrane-bound FtsN_cyto_His and FtsQ_cyto_His. This co-migration arises via their common binding partner FtsA.

### FtsZ, FtsA and FtsN self-organize via different underlying dynamics

In our experiments, FtsN_cyto_His, FtsA and FtsZ formed similar dynamic structures on the membrane. To better understand the underlying dynamics of protein patterns formation and the mechanism of peptide co-migration, we analyzed the exchange of proteins in fluorescence recovery after photobleaching (FRAP) experiments (Fig. 2g and Supplementary Fig. 4). All proteins showed rapid turnover in these assays. However, the time-scales of fluorescence recovery for FtsZ, FtsA and FtsN_cyto_His were very different. FtsZ showed the fastest recovery with a characteristic recovery time of 7.5 ± 3.5 s (n=10), while the pattern of FtsA and FtsN_cyto_His recovered about 8- and 3-fold slower (FtsA: □_05_= 64 ± 15 s, (n = 3), FtsN_cyto_His: L_05_= 27.61 ± 10 s (n = 10), Supplementary Fig. 4b, Supplementary Video 7). To discriminate between the mechanisms of protein recovery, we analyzed how the intensity profile across the bleached area changed over time. Consistent with lateral diffusion contributing to the recovery of FtsA and FtsN_cyto_His, the edges of their intensity profiles smeared out over time (Fig. 2g). In contrast, FtsZ recovered by exchanging proteins with the soluble pool in the buffer, as the shape of the intensity profile did not change (Fig. 2g) Therefore, these experiments show that although all three proteins co-migrate on the plane of the membrane, their underlying dynamics are fundamentally different.

### FtsN follows FtsZ/FtsA co-filaments via a diffusion-and-capture mechanism

To characterize the behavior of the proteins in greater detail we performed single-molecule experiments. By adding small amounts of fluorescently labeled proteins, we were able to monitor the dynamics of all three proteins during pattern formation (Fig. 3a, Supplementary Video 8). While FtsZ particles were stationary, FtsA molecules showed slow diffusion and the behavior of FtsN_cyto_His peptides could be divided in two populations with slow and fast diffusion, respectively (Fig. 3b). For all proteins the velocity autocorrelation was zero, indicating uncorrelated behavior of the individual protein molecules, which is inconsistent with directed motion (Fig. 3c).

**Figure 3.**
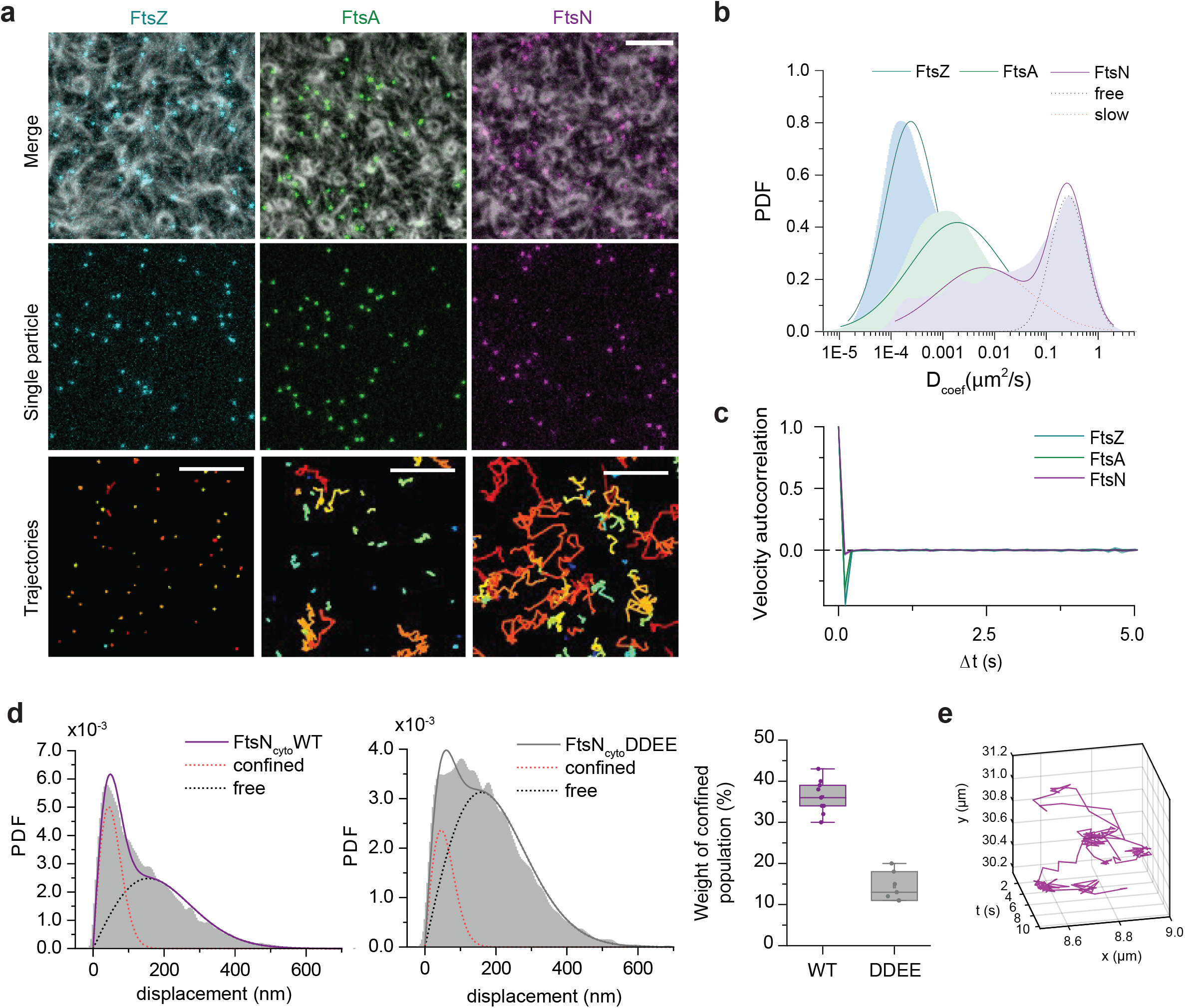
FtsN_cyto_His follows moving FtsA/FtsZ cofilaments via a diffusion-and-capture mechanism. a. Single-molecule imaging reveals different diffusive behavior of the proteins. Top rows: Small amounts of Cy5-labelled molecules are added to follow individual FtsZ (cyan), FtsA (green) and FtsN_cyto_His (magenta) particles in the presence of an Alexa488-FtsZ pattern. Scale bars are 5 μm. Bottom row: Corresponding trajectories showing different diffusive behavior of FtsZ, FtsA, and FtsN_cyto_His. Scale bars are 2 μm. Supplementary Video 8.
b. Histograms of diffusion coefficients (D_coef_) of single trajectories for FtsZ, FtsA and FtsN_cyto_His. The probability density distribution for FtsZ can be fitted to a single Gaussian (cyan) with a mean diffusion constant below 1 × 10^−4^μm^2^/s, which corresponds to immobile molecules. FtsA has an average D_coef_ of 0.0016 ± 0.0003 μm^2^/s (n = 3) approximated with a single Gaussian (green). Two populations can be observed for FtsN_cyto_His: fast diffusing peptides (black dotted line) and slow diffusing peptides (red dotted line) (n = 4).
c. On the single-molecule level FtsZ, FtsA and FtsN_cyto_His do not move directionally in contrast to the ensemble level. The velocity autocorrelations of single-molecule trajectories (averaged between independent experiments, n = 3) fluctuates around 0.
d. Step-size distribution of fast diffusing peptides for FtsN_cyto_His (left) and FtsN_cyto-D_DEEHis (middle). The weighted contribution of the confined population is 2-fold higher for FtsN_cyto_His in comparison to FtsN_cyto-D_DEEHis (right).
e. Example for a trajectory of a single FtsN_cyto_His molecule. The peptide is captured by FtsA/FtsZ filaments and stays confined in this region. An average confinement time of 0.41 ± 0.02 s for FtsN_cyto_His (combined data from independent experiments, n=3 experiments, total 4702 trajectories) was estimated from single exponential fits (R^2^ =0.98). Supplementary Video 9.

How can FtsNcytoHis co-migrate with treadmilling FtsZ/FtsA cofilaments if single molecules do not show directed motion? To answer this question, we further analyzed the behavior of single molecules of FtsN_cyto_His and compared it to FtsN_cyto-DDEE_His, which showed only weak overlap with FtsZ/FtsA cofilaments (Fig.1h and Supplementary Fig. 2). Compared to the wild-type peptide, FtsN_cyto-DDEE_His showed a larger fast-diffusing population of particles and a smaller population with slow diffusion. This is consistent with FtsA capturing single peptides of FtsN diffusing on the membrane. A closer inspection of the trajectories of the slow-diffusing populations for mutant and wild-type FtsN_cyto_His revealed that both data sets contained trajectories with confined diffusion as well as completely immobile molecules (Supplementary Fig. 5a-b). To exclude any contribution of stationary, non-specifically immobilized peptides, we considered only the fast-moving population for our further analysis. Calculating a histogram of displacement lengths, we identified two subpopulations of the fast-diffusing peptides that differed in their distribution of step sizes (Fig. 3d). For FtsN_cyto_His, the weight of the population showing short displacements was about 3 times higher than for FtsN_cyto-DDEE_His suggesting a transient capturing due to specific interactions between FtsN_cyto_His, involving the amino acid stretch RRKK, and FtsA/FtsZ filaments. Indeed, most trajectories of single FtsN_cyto_His molecules included short periods with confined motion (Fig. 3e) most likely arising from transient capturing events.

To quantify these capturing events, we identified clusters of short displacements within peptide trajectories in the presence of FtsZ and FtsA. We calculated an average duration of confinement of 0.41 ± 0.02 s for FtsN_cyto_His (n = 3 experiments with 4702 trajectories) and a slightly lower value for the DDEE peptide (0.36 ± 0 02 s, n = 3 experiments with 1763 trajectories). However, the frequency of confinement events was 3-fold higher for wildtype FtsN_cyto_His than for the DDEE mutant (Fig S5e-g, Supplementary Video 9) consistent with the decreased colocalization of the mutant with FtsA/FtsZ filaments on the membrane.

Although this approach cannot precisely determine interaction kinetics due to photobleaching and a possible contribution of molecular crowding to peptide confinement, the analysis shows that FtsN_cyto_His remained at FtsA/FtsZ co-filament bundles for an about 10-20-fold shorter time than FtsZ monomers stayed on the membrane^2,19^ and that wildtype FtsN_cyto_His is trapped more often than FtsN_cyto-D_DEEHis. Thus, our single-molecule analysis, in combination with MST (Fig. 1i) and FRAP (Fig. 2g) experiments, reveals that FtsNcytoHis is rapidly binding and unbinding to the treadmilling scaffold provided by the FtsZ/FtsA co-filament on the membrane.

To conclude, these results show that FtsNcytoHis is organized by FtsZ/FtsA co-filaments following a diffusion-and-capture mechanism and that the observed migratory behavior of the three proteins corresponds to an emergent property of this self-organized system: while on the ensemble level all proteins move directionally (Fig. 2e), the individual molecules show uncorrelated diffusive or stationary behavior (Fig. 3c).

### FtsN increases the persistency of the FtsZ cytoskeleton

FtsN has been shown to promote the stability of the divisome^20^. However, *in vivo* experiments could not provide evidence on how this stabilization might work. We realized that large-scale structures of FtsZ filaments are highly dynamic, but more persistent in the presence of FtsN_cyto_His (Fig. 4a, dashed square). To quantify the degree of reorganization, we performed an autocorrelation analysis, comparing the similarity of the pattern as a function of the time lag Δt. In the case of FtsA and FtsZ alone, we observed a fast decay (*τ* = 94.8 ± 28.5 s, n = 9) in the autocorrelation function. With FtsN_cyto_His, this decay was slowed down almost 3-fold (*τ* = 258 ± 43s, n = 9). This increased persistence of the pattern could not be observed for FtsNcyto-DDEE His (Fig. 4b).

**Figure 4.**
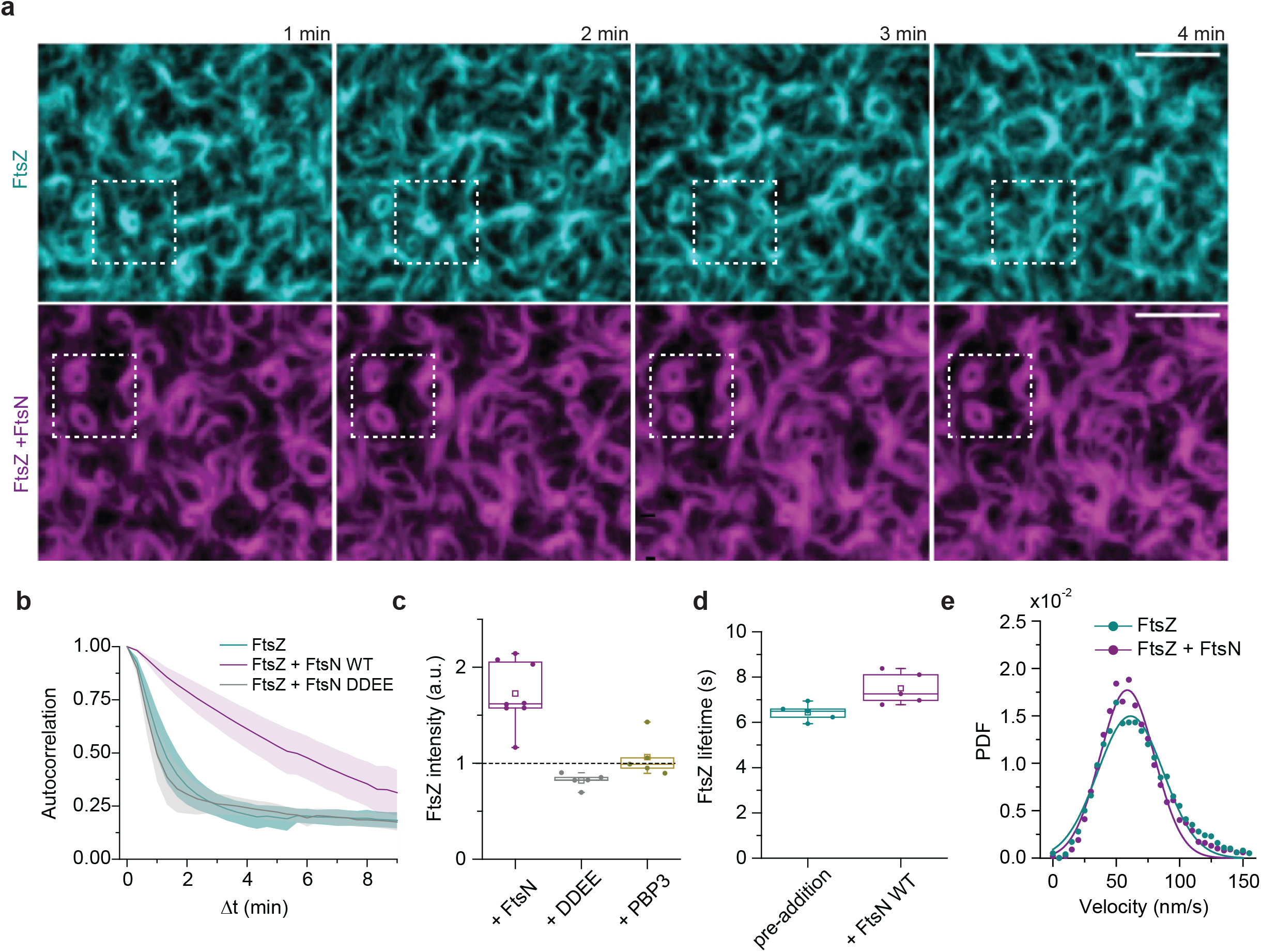
FtsN_cyto_His feedback to FtsZ leads to a higher spatial persistency. a. Montages of representative micrographs from two time-lapse movies. In the absence of FtsN_cyto_His, the pattern formed by FtsA and Cy5-FtsZ (cyan, top) constantly reorganizes in the plane of the membrane. The white dashed squares highlight an FtsZ ring that forms within 1 min and re-arranges within the next 4 min. After addition of FtsN_cyto_His (magenta, bottom), the cytoskeletal pattern of Cy5-FtsZ becomes more persistent in time and spatially localized: The two rings of FtsZ within the white dashed squares can be observed for much longer. Scale bars are 5 μm.
b. Autocorrelation analysis of the spatial FtsZ pattern (cyan, n = 9) is consistent with rapid reorganization. In the presence of FtsN_cyto_His (magenta, n = 9) the pattern is more persistent, as reflected in a slower decay in the spatial autocorrelation over lag-time. In contrast, addition of FtsN_cyto-DDEE_His (grey, n = 5) does not affect pattern reorganization.
c. In a continuous time-lapse experiment, addition of FtsNcytoHis leads to a 1.8-fold increase in FtsZ intensity (p-value = 1 × 10^−4^). No significant change was measured after addition of FtsN_cyto-DDEE_His or PBP3_cyto_His. The total intensity of FtsZ after peptide addition was normalized by the FtsZ pattern intensity before their addition.
d. The life-time of FtsZ monomers on the membrane is increased from 6.6 ± 0.3 s (cyan) to 7.7 ± 0.6 s (magenta) upon addition of FtsN_cyto_His (p-value = 0.02),
e. At the same time, FtsZ treadmilling velocity is not significantly changed (p-value = 0.26). Gaussian fits to velocity histograms (solid lines) resulted proposed in velocity values of v = 61.2 ± 25.9 nm/s for FtsZ alone and v = 58.7 ± 21.5 nm/s for FtsZ after FtsNcytoHis addition.

What could be the mechanism for the increased persistence? Comparing the fluorescence intensity of FtsZ before and after addition of the peptides, we found a 2-fold increase for FtsN_cyto_His, but not for FtsN_cyto-DDEE_His or PBP3_cyto_His (Fig. 4c). Furthermore, the mean lifetime of FtsZ monomers on the membrane slightly increased after FtsN_cyto_His addition, while the FtsZ treadmilling velocity remained unchanged (Fig 4d-e). As the direct binding partner of FtsN is FtsA, we also measured the recovery half-time of FtsA by FRAP experiments and found that FtsN_cyto_His decreases the turnover of FtsA by 2-fold (Supplementary Fig. 6a-c). To conclude, our data shows that FtsN_cyto_His is able to stabilize the FtsZ pattern, presumably by slowing down the turnover of FtsA and providing more binding sites for FtsZ filaments on the membrane.

## Discussion

In this work, we have reconstituted part of the *E. coli* cell division machinery to study how the Z-ring organizes transmembrane proteins that act on the opposite side of the cell membrane. We found that the membrane-bound cytoplasmic tails of FtsN and FtsQ co-migrate with treadmilling FtsZ/FtsA co-filaments. Our findings support a diffusion-and-capture mechanism, in which filaments of FtsZ and FtsA dynamically recruit FtsN and FtsQ diffusing in the membrane. As the treadmilling filament moves forward, these proteins follow the effective translational motion of the filament on the plane of the membrane (Fig. 5). At the depolymerizing end of the treadmilling FtsZ polymer, the accumulation of FtsN and FtsQ will disperse and bind to the same or another treadmilling filament. Importantly, individual protein complexes are stationary during treadmilling, suggesting that although proteins migrate collectively, the components of the divisome do not move.

**Figure 5:**
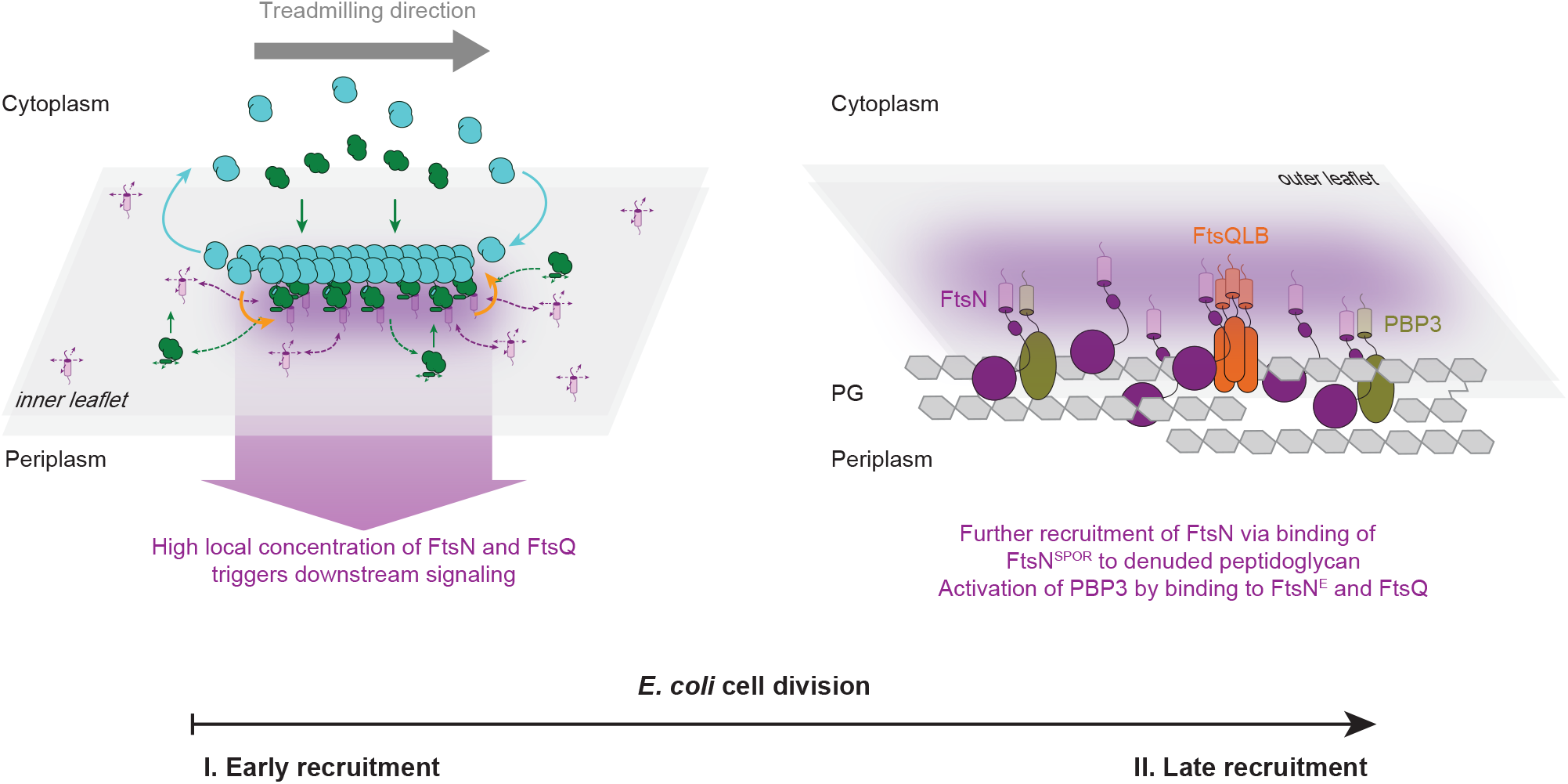
A model of FtsN co-migration with the treadmilling FtsZ cytoskeleton Left: Schematic illustration of the mechanism of co-migration based on our results. The treadmilling FtsZ/FtsA co-filament acts as a dynamic scaffold that recruits transmembrane proteins diffusing in the bilayer. As the filament moves forward, it recruits new proteins, while they rapidly disperse at the depolymerizing end. Due to the local increase in concentration, weak interactions between proteins can be sufficient to dynamically assemble the divisome. The positive feedback provided by FtsN is shown as yellow arrows. **Right**: At later stages, FtsN can further accumulate at the cell division site due to its binding to other periplasmic proteins and to the septal peptidoglycan layer, which has been modified by cell division specific amidases.

Despite the strong colocalization of their fluorescence signals, the corresponding complexes form only transiently, with an estimated average lifetime on the millisecond scale, about 10-20 times shorter than the average residence time of FtsZ monomers on the membrane. Accordingly, the co-migration of proteins results from weak, highly dynamic interactions. Compared to static interactions, this could have important advantages for the cell as it would allow the Z-ring to initiate divisome formation from freely diffusing components, without the need to disassemble stable complexes, and divisome assembly could quickly respond to repositioning and constriction of the Z-ring. In addition, this mechanism would avoid local complex clustering at the septum, facilitating homogeneous synthesis of the cell pole.

Our data suggests that FtsZ/FtsA cofilaments function as a binding platform that locally increases the density of downstream interaction partners above a critical threshold to promote weak interactions to other transmembrane proteins of the divisome. This dynamic enhancement of protein-protein interactions could increase the precision of cell division, as individual freely diffusing molecules would not be able to trigger a downstream response outside of this dynamic accumulation. Importantly, this mechanism could also promote directionality of PG synthesis by establishing a narrow zone of signaling activity that moves around the cell diameter at the nascent division site. However, cell division proteins activated in this zone would be stationary, which would be in contrast to cell wall growth during cell elongation, where the proteins involved move during PG synthesis^21–23^. As we did not observe interaction between FtsA and PBP3_cyto_His, the observed directed motion of individual PBP3 molecules *in vivo*^4^ might originate from a more indirect mechanism, presumably involving additional interactions with other cell division proteins such as FtsW or FtsN either in the membrane or in the periplasm ^24,25^.

Although FtsN has long been thought to be the last protein to appear at the division site^26^, it is now evident that its early interaction with FtsA is important for divisome assembly^10^. Accordingly, FtsN is recruited to the division site by two consecutive, but independent signals. At an early stage of divisome assembly, which we have reconstituted in this work, FtsN is recruited to midcell via the interaction of its cytoplasmic tail with FtsA in the Z-ring^7,10,14,18^ After the onset of septation during synthesis of PG, FtsN binds via its periplasmic SPOR domain to PG degraded by division specific amidases, which leads to further accumulation of FtsN at the division site^27^. At the same time, FtsN interacts with the PG synthases PBP1B and PBP3, and stimulates the former *in vitro*^25,28^. This is reminiscent of the situation in S. *aureus*, where the divisome is first assembled in an FtsZ treadmilling-dependent step, before PG synthesis provides the contractile force for complete septum formation^29^.

The FtsZ-FtsA-FtsN/Q system is an example of a reaction-diffusion process with emergent properties in which individual components show different behaviours at single molecule level compared to an ensemble. How does this protein system compare to other mechanisms that the cell uses to organize its interior? The best studied example for proteins able to hitchhike on a polymerizing filament are probably +TIP tracking proteins^30^, which follow the growing microtubule ‘plus’ end while individual molecules are continuously turning over. Similarly, actin binding proteins were suggested to co-migrate with treadmilling actin filaments from the neuronal cell body to the axon tips^31^. These examples suggest that co-migration of proteins with dynamics filaments by transient binding is a widely conserved mechanism used for the spatial organization of the cell. Therefore, we believe that this work not only sheds light on how bacterial cells divide but more generally highlights an important mechanism to organize and transmit spatiotemporal information in the living cell.

## Supporting information

## Acknowledgements

We acknowledge Paulo Caldas for help with single molecule lifetime analysis and all other members of the Loose lab at IST Austria for helpful discussions, Mercedes Jimenez, Ana Raso and Noelia Ropero for providing AlexaFluor488- and AlexaFluor647-labelled FtsA for MST and AUC experiments. We want to thank Ralf P Richter (Leeds University, UK) for providing DODA-tris-NTA phospholipids as well as Jacob Piehler and Christian Richter (University of Osnabrück, Germany) for the SLIMfast single molecule tracking software and help with the confinement analysis. We thank Jeff Errington and Heath Murray (both Newcastle University, UK) for critical reading of the manuscript and Jan Brugués (MPI-CBG and MPI-PKS, Dresden, Germany) for help with MATLAB programming and reading the manuscript. This work was supported by the European Research Council through grant ERC-2015-StG-679239 to ML, HFSP LT 000824/2016-L4 and EMBO ALTF 1163-2015 to NB, a grant from the Ministry of Economy and Competitiveness of the Spanish Government BFU2016-75471-C2-1-P to CA and GR, and a Wellcome Trust Senior Investigator award (101824/Z/13/Z) to WV.

## Author contributions

Conceptualization, N.B., V.H.-R., W.V. and M.L.; Methodology, N.B., V.H.-R., C.A., M. L.-P., G.R. W.V. and M.L.; Software, M.L.; Validation, N.B. and P.R.; Formal Analysis, N.B., P.R., V.H.-R., C.A. and G.R.; Investigation N.B., P.R. and V.H.-R.; Resources N.B., P.R., V.H.-R., C.A., M. L.-P., G.R. and M.L.; Data Curation, N.B., P.R. and M.L.; Writing - Original Draft, N.B., C.A., G.R. and M.L.; Writing - Review & Editing, N.B., P.R., V.H.-R., W.V. and M.L.; Visualization, N.B. and P.R.; Supervision, N.B., W.V. and M.L.; Project Administration, W.V. and M.L.; Funding Acquisition, N. B., W.V. and M.L..

## Material & Methods

### Reagents

2 Reagents
| ***Strain, Reagent or Resource*** | ***Source*** | ***Identifier*** |
| --- | --- | --- |
| ***E. coli strains*** |  |  |
| BL21 (DE3) |  | C2527 |
| C41(DE3) | Lucigen | 60442 |
| DH5α | | 18265017 |
| ***Chemicals*** |  |  |
| cOmplete™, EDTA-free Protease Inhibitor Cocktail | Roche Molecular Biochemicals | 5056489001 |
| DNase | ThermoFisher Scientific | 90083 |
| Glycerol | Sigma-Aldrich | G5516 |
| IPTG | Bartelt | 6.259 683 |
| Imidazole | Sigma-Aldrich | I2399 |
| Piperazine-N,N-bis(2-ethanesulphonic acid) (PIPES) | Sigma-Aldrich | P6757 |
| Hydrogen Peroxide H<sub>2</sub>O<sub>2</sub> | Sigma-Aldrich | 216763 |
| Sulfuric acid H<sub>2</sub>SO<sub>4</sub> 95-97% | Sigma-Aldrich | 1007311000 |
| β-mercaptoethanol | Sigma-Aldrich | M6250 |
| Lysozyme | Sigma-Aldrich | 62971-10G-F |
| HisPur™ Ni-NTA resin | ThermoFisher Scientific | 88221 |
| 7M Sortase | Addgene | Plasmid #51141 |
| Tris(2-carboxyethyl)phosphine hydrochloride (TCEP) | Sigma-Aldrich | C4706 |
| Protino Ni-TED resin | Lactan (Macherey Nagel) | 74520030 |
| Vivaspin 20 centrifugal concentrators 5-kDa | Sigma-Aldrich | Z614599-48EA |
| Catalase | Sigma Aldrich | C40 |
| GTP | ThermoFisher Scientific | R0461 |
| ATP | ThermoFisher Scientific | R0441 |
| Trolox (6-Hydroxy-2,5,7,8-tetramethylchromane-2- | Sigma Aldrich | 238813 |
| carboxylic acid) |  |  |
| Glucose Oxidase | AL-Labortechnik<br>(SERVA) | 22778.01 |
| Phenylmethylsulfonylfluoride<br>(PMSF) | Sigma-Aldrich | P7626 |
| n-Dodecyl-β-D-Maltopyranoside (DDM) | Anatrace | D310 |
| **Lipids** |  |  |
| **DOPC (1,2 dioleoyl-sn-glycero-3-phosphocholine)** | Avanti Polar Lipids | 850375C |
| **DOPG (1,2-dioleoyl-sn-glycero-3-phospho-(1'-rac-glycerol))** | Avanti Polar Lipids | 840475C |
| DGS Ni-NTA(1,2-dioleoyl-sn-glycero-3-[(N-(5-amino-1-carboxypentyl)iminodiacetic acid)succinyl] (nickel salt)) | Avanti Polar Lipids | 790404C |
| Cholesterol | Sigma Aldrich | C8667 |
| **Tris-DODA-NTA** | Ralf Richter Lab | N/A |
| **Peptides** |  |  |
| FtsL<sub>cyto</sub>His | Biomatik | Supplementary table 1 |
| FtsN<sub>cyto-DDEE</sub>His | Biomatik | Supplementary table 1 |
| FtsN<sub>cyto-RAAK</sub>His | Biomatik | Supplementary table 1 |
| FtsN<sub>cyto-D5N</sub>His | Biomatik | Supplementary table 1 |
| PBP3<sub>cyto</sub>His | Biomatik | Supplementary table 1 |
| FtsQ<sub>cyto</sub>His | Biomatik | Supplementary table 1 |
| CLPETG | Biomatik | N/A |
| TMR-LPETGG | GenScript | N/A |
| **Other** |  |  |
| Pierce™ BCA Protein Assay Kit | ThermoFisher Scientific | 23227 |
| Pierce™ Coomassie (Bradford) Protein Assay Kit | ThermoFisher Scientific | 23200 |
| PD10 desalting column | Sigma Aldrich | GE17-0851-01I |
| Precision cover glasses thickness No. 1.5H, 24 x 50 | VWR (Marienfeld) | 107222 |
| mm |  |  |
| Norland Optical Adhesive 63, 10Z. | APM Technica | 284239 |
| HisTrap HP column | GE Healthcare | 17524802 |
| HiTrap desalting column | GE Healthcare | 17140801 |
| **Dyes** |  |  |
| Sulfo-Cyanine-5-maleimide | Lumiprobe | 23380 |
| Alexa Fluor® 488 C5 Maleimide | ThermoFisher Scientific | A10254 |
| CF 488 | Sigma-Aldrich | SCJ4600016 |
| Tetramethylrhodamine |  |  |
| AlexaFluor647-NHS ester | ThermoFisher Scientific | A37573 |
| AlexaFluor488-NHS ester | ThermoFisher Scientific | A20000 |
| **Software and algorithms** |  |  |
| Fiji | <sup>1</sup> |  |
| Matlab | MathWorks | https://fiji.sc |
| Origin | Origin (OriginLab, Northampton, MA) | https://www.mathworks.com |
| **Deposited data** |  | http://www.originlab.com |
| Velocities |  |  |
| Correlation Coefficient |  |  |
| FRAP recovery |  |  |
| FtsZ residence time |  |  |
| Autocorrelation |  |  |

### Protein biochemistry Purification and fluorescence labelling of FtsZ

The gene coding for FtsZ (UP P0A9A6) was cloned into a modified pML45 vector containing an N terminal His_6_-SUMO fusion protein plus seven residues (AEGCGEL) for maleimide-coupling of thiol-reactive dyes and further fluorescence detection (hereafter pML45-GCG-FtsZ vector)^2^. FtsZ was expressed in *Escherichia coli* C41 (DE3) cells, which were grown at 37°C in Terrific Broth (TB) medium supplemented with 100μg/ml ampicillin. Cultures were induced at an OD_600_ of 0.8 with 1mM isopropyl-βthiogalactopyranoside (IPTG) and incubated for 5h at 37°C.

Purification of FtsZ was performed as follows. After centrifugation, the pellet was resuspended in buffer A (50 mM Tris-HCl pH□8.0, 500 mM KCl, 2□mM β-mercaptoethanol, 10% glycerol) plus 20 mM Imidazole and supplemented with EDTA-free protease inhibitor cocktail tablets (Roche Diagnostics). The cell pellet was frozen and stored at −80□°C until further processing. Cells were lysed using a cell disrupter (Constant Systems Cell TS 1.1) at a pressure of 1.36Kbar. The resulting lysate was incubated with 2.5 mM MgCl2 and 1 mg/ml DNase for 15 min, and the cell debris was removed by centrifugation at 60,000xg for 30 min at 4°C. The clarified lysate was incubated with nickel-nitrilotriacetic (Ni-NTA) resin (HisPur^TN^ Ni-NTA resin, ThermoFisher Scientific) for 1 h at 4□°C. The resin was washed with buffer A containing 10 mM imidazole and 20 mM imidazole. The fusion protein was eluted with buffer A containing 250 mM imidazole. Subsequently, peak FtsZ frations were dialyzed overnight at 4°C against buffer B (50□mM Tris-HCl pH□8.0, 300□mM KCl, 10% glycerol) in the presence of His_6_-tagged SUMO protease (Ulp1) at a protease:sample molar ratio of 1:100. This cleavage left the peptide AEGCGEL at the N terminus of the protein. The digested sample was passed several times through Ni-NTA resin (HisPur^TN^ Ni-NTA resin, ThermoFisher Scientific) previously equilibrated with buffer B to remove His_6_-containing molecules. The flow-through was collected, and polymerization-competent FtsZ was enriched by CaCl_2_-mediated polymerization. For this, the sample buffer was exchanged into buffer C (50 mM piperazine-N,N-bis(2-ethanesulphonic acid) (PIPES) pH 6.7, 10 mM MgCl_2_) using a PD10 desalting column and the sample was warmed to room temperature. CaCl_2_ and GTP were added to final concentrations of 10□mM and 5□mM respectively After 20 min of incubation, the mixture was centrifuged at 15,000 × g for 2□min to pellet polymeric FtsZ, which was then resuspended in buffer D (50 mM Tris-HCl pH 7.4, 50 mM KCl, 1 mM EDTA, 10% glycerol).

Protein identity and purity were assessed by 12.5% Glycine-SDS-PAGE stained with Coomassie blue. Protein was concentrated by ultrafiltration with Vivaspin 20 filter devices of 5-KDa cut-off (Sartorius) and protein concentration was determined by Bradford assay.

For specific labeling of FtsZ, thiol-reactive dyes Alexa Fluor^®^ 488 C5 Maleimide (ThermoFisher Scientific) or Sulfo-cyanine 5-maleimide (Lumiprobe) were dissolved in DMSO according to the manufacturer’s instructions. FtsZ was incubated first with 100×molar excess of Tris(2-carboxyethyl)phosphine hydrochloride (TCEP) for 20 min at room temperature and then with 10× molar excess of the thiol-reactive dye Alexa Fluor^®^ 488 C5 Maleimide or Sulfo-cyanine 5-maleimide. Immediately after, the reaction was extensively dialyzed against buffer D overnight at 4°C and loaded on a PD10 desalting column to remove CaCl_2_, GTP and free dye. Peak fractions were collected and after determining the protein concentration, FtsZ was aliquoted, flash frozen in liquid nitrogen and stored at −80 °C.

### Purification and fluorescence labelling of FtsA

The gene coding for FtsA (UP P0ABH0) was cloned into a modified pTB146 vector as described elsewhere^2^. The resulting vector, pML60, contains an N terminal His_6_-SUMO fusion protein plus a pentaglycine tag for fluorescent labeling. FtsA was expressed in *E. coli* C41 (DE3) cells, which were grown at 37 °C in 2xYT medium supplemented with 100μg/ml ampicillin. Cultures were induced at an OD_600_ of 0.6 with 1mM IPTG and incubated overnight at 18 °C.

Cells were harvested by centrifugation (5,000 × g, 30 min at 4□°C), frozen in liquid nitrogen and stored at −80 °C until further use. The pellet was resuspended in buffer A (50 mM Tris pH 8.0, 500 mM KCl, 10 mM MgCl_2_) supplemented with 1 mg/ml lysozyme, EDTA-free protease inhibitor cocktail tablets, 1mg/ml DNase I, 0.5 mM DTT and 0.5 mM ADP (DTT and ADP were freshly added). Cells were lysed using a cell disrupter at a pressure of 1.36 kbar. Cell debris was removed by two successive centrifugation steps at 24,000 × g for 1 h and 35,000 × g for 30 min at 4 °C. The clarified lysate was incubated with nickel-(tris-carboxymethyl ethylene diamine) resin (Protino Ni-TED, Macherey Nagel) for 1 h at 4°C. The resin was washed with buffer A and buffer A supplemented with 5 mM imidazole. The fusion protein was eluted using buffer A with 250 mM imidazole. Immediately after the affinity purification buffer A was exchanged to buffer B (50 mM Hepes, pH 7.5, 500 mM KCl, 10 mM MgCl_2_, 0.5 mM ADP, 20% glycerol) using a PD10 column, and the protein was stored at 4°C overnight. The sample was incubated at 30°C for 1.5 h in the presence of Ulp1 at a protease:sample molar ratio of 1:100. The cleaved sample was passed several times through Protino Ni-TED resin, previously equilibrated with buffer B to remove His6-containing molecules and FtsA was collected in the flow-through. Protein identity and purity were assessed by 12.5% glycine/SDS-PAGE stained with Coomassie Blue. As described above for FtsZ, protein was concentrated by ultrafiltration, protein purity was assessed by 12.5% Glycine-SDS-PAGE,and the protein concentration was determined by Bradford assay.

For TIRF microscopy and single particle tracking, FtsA was labeled with fluorescent dyes using sortagging (Theile *et al*. 2013), where a pentaglycine at the N-terminus of FtsA was conjugated to the TMR-dye labelled peptide by Sortase 7M from S. *aureus* (Addgene plasmid number #51141). The reaction was performed by mixing 0.5 −1 mM LPETG/A-containing probe (from 5 mM stock in DMSO) with 10 μM target protein and 10 μM Sortase 7M during overnight dialysis into buffer B at 4°C. The protein was further purified by size-exclusion chromatography on a HiLoad Superdex 200 10/300, previously equilibrated with buffer B (supplemented with 10% of glycerol) during which free labeled peptide and His_6_-sortase were removed. The final protein corresponding to labelled monomeric FtsA was collected.

For MST experiments, FtsA was purified as described before^2^, but with 10 mM CHAPS in the elution buffer. The detergent was later removed using a Hi-Trap Desalting column, before storing the protein in storage buffer. The protein was labelled with amine-reactive Alexa647-NHS ester or Alexa488-NHS ester following the instructions by the manufacturer. Briefly, FtsA was equilibrated in 50 mM Hepes, 500 mM KCL, 10mM MgCl2, 10% glycerol, 1mM ADP at pH 7.5 and then mixed with the probe in DMSO at a 1:1 protein:probe molar ratio for 30 min on ice with stirring. Unbound probe was removed by desalting into the same buffer using a 5 ml HiTrap desalting column. FtsA concentration was quantified by Bradford Assay and dye concentrations were determined by absorbance measurements at 690 and 490 nm for AlexaFluor647 and AlexaFluor488, respectively. The degree of labelling was below 0.9 mol fluorophore/mol protein in all cases.

### Purification of full length FtsN

Full-length FtsN modified with a C-terminal oligohistidine tag was expressed in *E. coli* BL21(DE3) pFE42 (ref. 3) and purified as described previously^4^ with minor modifications. Briefly, overproduction was induced by the addition of 1 mM IPTG and cells were incubated for 2 h at 37°C. Cells were harvested by centrifugation (7000×g, 15 min, 4°C), and then resuspended in buffer FtsN-1 (25 mM sodium phosphate, 1 M NaCl, pH 6.0) in the presence of protease inhibitor cocktail (Sigma, USA) and 100 μM phenylmethylsulfonylfluoride (PMSF). After disruption by sonication (Branson Digital, USA), the sample was centrifuged (130 000×g, 1 h, 4°C) and the resulting membrane pellet was resuspended in buffer FtsN-2 (25 mM sodium phosphate, 1 M NaCl, 40 mM imidazole, 1% dodecyl maltoside (DDM), pH 6.0) and incubated overnight at 4°C. After centrifugation (130 000×g, 1 h, 4°C), the supernatant was applied to a 5 ml HisTrap HP column using an FPLC system. The column was washed with four column volumes of buffer FtsN-3 (25 mM sodium phosphate, 1 M NaCl, 40 mM imidazole, 0.05% DDM, pH 6.0). Bound protein was eluted step-wise with buffer FtsN-4 (25 mM sodium phosphate, 1 M NaCl, 400 mM imidazole, 0.05% DDM, pH 6.0). FtsN-His was dialyzed into buffer FtsN-5 (25 mM sodium phosphate, 500 mM NaCl, 0.05% DDM, 10% glycerol, pH 6.0) and stored in aliquots at −80°C.

### Peptide reconstitution and labeling

The cytoplasmic peptides of FtsN, PBP3, FtsL and FtsQ as well as the FtsN mutants D5N, DDEE and RAAK, containing a 6-His Tag and an N-terminal cysteine residue were purchased from Biomatik (Supplementary Table 1).

For labelling the lyophilized peptides were dissolved in buffer A (150 mM KCl, 50 mM HEPES, 10% glycerol, pH 7.4) to 10 mg/ml. Prior to the labeling step, a 20x molar excess of TCEP was added and incubated for 20 min at ambient temperature. Afterwards a 10x molar excess of thiol-reactive maleimide dyes was added and the sample was incubated either overnight at 4□°C or for 4 h at ambient temperature. The free dye was removed by affinity purification using Ni-NTA resin. The beads were washed extensively with 100 column volumes of buffer A, 100 column volumes of washing buffer A containing 10 mM Imidazole and eluted with 850 μL of buffer A containing 250 mM Imidazole. The peptides were dialyzed into storage buffer B (150 mM KCl, 20 mM HEPES, 10% glycerol, pH 6.0). The concentration was determined using the Pierce BCA Protein assay kit. Samples were aliquoted and stored at −80□°C, and used within 4 weeks.

### Microscale thermophoresis

For MST experiments, 25 nM FtsA labelled with AlexaFluor647 was titrated with increasing concentrations of unlabeled full-length FtsN, PBP3_cyto_His, FtsQ_cyto_His, FtsL_cyto_His or FtsN_cyto_His. Titrations with full-length FtsN were equilibrated in 10 mM Hepes pH 7.4, 2 mM MgCl_2_, 0.2 mM TCEP, 0.19% DDM and either 100 or 300 mM KCl. Titrations with the cytoplasmic tail peptides were equilibrated in 50 mM Tris/HCl pH 7.4, 150 mM KCl, 5 mM MgCl_2_ and 0.2 mM TCEP. After 10 min incubation at room temperature, samples were loaded in premium coated capillary tubes (NanoTemper, Germany) and measured in a Monolith NT.115 (NanoTemper, Germany) equipped with a blue and a red filter set. Data were acquired using 60% MST and 50% LED settings. Red fluorescence was measured for 3 s before applying a thermal gradient for 20 s.

Binding curves were obtained by plotting the normalized change in fluorescence intensity after the 20 s gradient against the concentration of titrated protein or peptide. Binding curves were then analyzed using a Hill Equation.

### Analytical ultracentrifugation

Sedimentation velocity (SV) experiments were performed to study the oligomerization of full length FtsN-His in the presence of detergent as well as its association with FtsA. For the latter, Alexa488-labelled FtsA was used in order to track changes in FtsA sedimentation upon addition of FtsN. Samples were thoroughly dialyzed in the indicated buffers before the SV experiments. FtsN on its own was loaded at 9-20 μM and equilibrated in buffer 10 mM Hepes pH 7.4, 2 mM MgCl_2_, 0.2 mM TCEP, 0.19% DDM and 100, 200 or 300 mM KCl. For interaction assays samples were equilibrated in the same buffer as above with 100 mM KCl, an FtsA-Alexa488 was loaded at 4 μM with FtsN present either at 1:1 or 1:4 FtsA:FtsN molar ratios. Experiments were carried out in a XL-I analytical ultracentrifuge (Beckman-Coulter Inc., USA) equipped with UV-visible and interference detection systems. Sedimentation profiles were registered by Raleigh interference and by absorbance at 280 nm in experiments with FtsN-His on its own or at 495 nm when FtsN was mixed with FtsA-AlexaFluor488. Sedimentation coefficient distributions were calculated by least-squares boundary modeling of sedimentation velocity data using the c(s) method as implemented in SEDFIT ^5^. Absorbance and Raleigh interference measurements were combined to study FtsN on its own solubilized in DDM to quantify the amount of bound DDM per molecule of FtsN ^6^

### Preparation of small unilamellar vesicles (SUVs) and supported lipid bilayers (SLB)

SUVs were prepared by rehydrating a lipid film composed of DOPC, DOPG and DOGS-Ni-NTA at a ratio of 66/33/1 mol-% or DOPC, DOPG (initial experiments) which mimics the lipid composition of the E. coli membrane (Vecchiarelli *et al.*, 2014).

To improve peptides attachment to membrane (critical for FRAP analysis and single particle tracking of FtsN_cyto_His) commercial mono-Ni-NTA lipid was replaced with DODA-tris-Ni-NTA at a ratio of 66.8/33/0.2 mol%.

Phospholipids dissolved in chloroform were mixed in a glass vial at the desired ratio and blown-dried with filtered N_2_ to form a thin homogeneous film. To remove residual chloroform the vials were kept under vacuum for 2-3 h. The lipid film was rehydrated in a swelling buffer (300 mM KCl, 50 mM Tris-HCl, pH 7.4) for 30 min at ambient temperature to yield a total lipid concentration of 5 mM. The mixture was vortexed rigorously and the resulting dispersion of multilamellar vesicles was repeatedly freeze-thawed (8-10x) in liquid N_2_. To obtain small unilamellar vesicles, the vesicle dispersion was tip-sonicated for 25 min on ice. Vesicles were centrifuged for 5 min at 10 000 ×*g* and the supernatant stored at −20° C to be used within two weeks.

Glass coverslips were cleaned in piranha solution (30% H_2_O_2_ mixed with concentrated H_2_SO_4_ at a 1:3 ratio) for 30 min, and extensively washed and sonicated for 10 min with double distilled H_2_O. Cleaned coverslips were stored for no longer than one week. Before each experiment, glass coverslips were dried with compressed air and cleaned in air plasma for 10 min. The reaction chamber was prepared by attaching a 0.5 ml Eppendorf tube, with its conical end removed, on a coverslip using ultraviolet glue (Norland optical adhesive 63), and exposing the chambers to UV light for 5 min.

Supported lipid membranes were formed by diluting the SUV dispersion in the reaction buffer (150 mM KCl, 50 mM Tris-HCl, 5 mM MgCl_2_, pH 7.4) to a lipid concentration of 0.5 mM. Formation of SLBs on the glass surface was initiated by addition of 4 mM CaCl_2_. After 30 min of incubation the surface were washed 10 times with an excess of reaction buffer to remove nonfused vesicles. The supported lipid membranes were used shortly after preparation.

### TIRF microscopy

All experiments were performed on a total internal reflection fluorescence (TIRF) microscope (“Till Photonics”) equipped with a 100x Olympus TIRF NA□1.49 DIC objective. The fluorophores were excited using stable laser lines at 488, 561 and 640 nm. The emitted fluorescence from the sample was filtered using an Andromeda quad-band bandpass filter (FF01-446-523-600-677). For dual-color experiments, an Andor TuCam beam splitter, equipped with a spectral long pass of 580 and 640nm and band pass filter of 525/50, 560/25, 710/80 (Semrock) was used. Time series were recorded using iXon Ultra 897 EMCCD Andor Cameras (X-8499 & X-8533) operating at 5 Hz frequency for standard acquisition and at 10 Hz for single particle tracking. The acquisition rate was varied based on the experiment, while exposure time was kept at 30-50 ms to minimize photobleaching.

The acquisition rate was varied based on the experiment, while exposure time was kept at 30-50 ms to minimize photobleaching.

### Peptide-filament colocalization assay

To prevent photobleaching during acquisition 30 mM D-glucose, 0.050 mg/ml glucose oxidase, 0.016 mg/ml catalase, 1-10 mM DTT and 1 mM Trolox were added to the reaction buffer, followed by the addition of nucleotides: 4 mM ATP and 4mM GTP. All components were diluted from 100x concentrated stock solutions stored at −20C (chemicals) and −80C (enzymes).

To allow FtsA and FtsZ to self-organize on a supported lipid membrane the reaction was started by adding a mixture of FtsA (0.45-0.6 μM) and FtsZ (1.5-1.8 μM) to 100 μL of the reaction volume in reaction buffer R(150 mM KCl, 50 mM Tris, 5 mM MgCl_2_, pH 7.4). Within this range of protein concentrations, we were able to observe dynamic patterns of treadmilling FtsZ filaments on the membrane. The protein system reached steady state dynamics, defined by a constant intensity of the proteins, in10 min min after the initiation of the reaction.

To perform co-localization experiments, we added 1 μM of fluorescently labeled, His-tagged peptides to the reaction at steady state, and recorded the experiment for another 20 min to quantify the degree of colocalization. FtsZ labeled with Alexa488, FtsA labeled with TMR and His_6_-tagged peptides (FtsN_cyto_His, FtsQ_cyto_His, FtsL_cyto_His, PBP3_cyto_His, FtsN_cyto-D5N_His, FtsN_cyto-RAAK_His and FtsN_cyto-DDEE_His) labeled with Cyanin-5 were used in colocalization experiments. For experiments with fluorescent FtsA and FtsN, unlabeled FtsZ and FtsA-TMR (30% of labeled FtsA) was used.

For peptide sorting experiments (Fig. 1 b,c), FtsN_cyto_His and FtsQ_cyto_His were attached to supported lipid membranes containing either 1 mol% Ni-NTA or 0.2 mol% tris-Ni-NTA to obtain a homogeneously distributed signal for several minutes with a 2 s/frame, 50 ms exposure time. Afterwards we added ATP, GTP, FtsA and labeled FtsZ to initiate formation of the protein pattern, and continued to acquire time-lapse movies for another 20 min. For FtsN_cyto_His and FtsQ_cyto_His colocalization experiments, peptides labeled with Alexa488 and Cyanin5 respectively were mixed in a molar ration 1:3 and attached to 0.5 mol% tris-Ni-NTA. The homogeneously distributed signal for both peptides was recorded for about 1 min before the addition of unlabeled FtsA and FtsZ, which triggered their organization into filament-like pattern. Time-lapse movies were acquired at 2 s/frames and 50 ms exposure time. To minimize cross-talk, the fluorescence signal of each channel was recorded sequentially.

To test the influence of dynamic reorganization of FtsZ filaments on the colocalization with FtsN_cyto_His, 4 mM GTP was replaced with 1 mM GMPCPP. Acquisition was started after the steady state was reached (10-15 min) and the rate was reduced to 5 s/frame to prevent excessive photobleaching.

### Image processing and analysis

For image processing, image stacks were imported using FIJI software^7^. For data analysis raw, unprocessed time-lapse movies were used. For visualization purposes and to remove noise, all micrographs shown in the manuscript were processed with the “Walking average”plugin of ImageJ averaging the signal of 4 consecutive frames.

### Colocalization analysis

For colocalization analysis, we first normalized the intensity of the time-lapse movie to correct for photobleaching and enhanced the contrast of both channels for visualization purposes. Next, movies were aligned with the “Linear Stack Alignment with SIFT” plugin prior to analysis to eliminate XY drift. To minimize the effect on non-specifically bound aggregates, regions of interest (20 x 20 μm) without protein aggregates were chosen for the colocalization analysis. The alignment of the two channels was adjusted with the 3TP align plugin (J. Anthony Parker, Beth Israel Deaconess Medical Center Boston). Finally we used the ImageJ plugin “Image CorrelationJ 1o” to quantify colocalization between the proteins and candidate peptides ^8^. This plugin measures the crosscorrelation coefficient (Pearson’s correlation coefficient, PCC) values between −1 and 1. In our experiments, we reached maximal PCC values of ~0.9, which indicates a high degree of colocalization, while the minimum value varied was 0.05, due to some residual signal of non-colocalized peptides diffusing on the membrane.

### Treadmilling velocity analysis

Initial analysis of the velocities for FtsZ, FtsA and FtsN was performed using kymographs: a line (width = 3 px) was drawn along the circumference of rings and corresponding kymographs were generated by the ImageJ function “Reslice”. The velocity (nm/s) was extracted from the resulting slope of intensity fluctuations.

For a more objective analysis and to obtain better statistics, we developed a novel automated protocol to quantify treadmilling velocities. First, intensity normalized movies were preprocessed with the “Linear Stack Alignment with SIFT” plugin prior to analysis, to eliminate XY drift. Next, we constructed intensity difference images of time lapse movies, by subtracting two frames separated by a given time delay from one another (i.e. I(x,y,t+Δt) - I(x,y,t)) using ImageJ’s “ImageCalculator” command. This resulted in a new time lapse movie of moving speckles corresponding to the amount of added protein at a given position (Supplementary Fig. 3). We chose time delays (Δt) that yielded optimal signal/noise ratios: 6 s for FtsZ, 14 s for FtsA and 12 s for FtsN. Next, we used TrackMate, a Fiji/ImageJ plugin for tracking of single particles 9 to quantify the treadmilling velocity of FtsZ, FtsA and FtsN. Speckles were detected using the LoG (Laplacian of Gaussian) detector from TrackMate with an estimated speckle diameter of 1 μm. In a first filtering step following the initial detection step, we took advantage of TrackMate’s quality criterion, assigning a quality value by summing the second order spatial derivatives of the Gaussian fit of the filtered image. The quality value is thus largest for bright speckle, with a diameter close to the specified value. We discarded 95% (97.5% in the case of FtsA) of all speckles to use only the ones of highest quality for further analysis. Additionally we only used speckles with a signal/noise ratio higher than 0.7 as calculated by TrackMate. These strict criteria are important to prevent tracking of non-correlated intensity fluctuations, which appear as random motion in the analysis. For trajectory building, we used the “Simple LAP tracker” function of TrackMate with the following criteria: Maximal linking distance = 0.5 μm; Maximal gap-closing distance = 1 μm; Maximal frame gap = 2. Furthermore, we only used trajectories longer than 6s for further analysis.

Trajectories were further analyzed using the MSDanalyzer toolbox in Matlab ^10^ to calculate the Mean-Squared-Displacement (MSD) and Velocity Autocorrelation. To obtain velocities from the MSD graph, we fitted a polynomial equation *MSD* = *v*^2^*t*^2^ * +4*Dt*, assuming a mixed behavior of single particles, namely directed motion and diffusion, taking into account only the first 7 frames: The first term represents the diffusive portion of speckle displacement, whereas the second term includes the velocity imparted by directed motion. In all cases, the value of the first term was negligible compared to the velocity of directed motion.

Velocity autocorrelation compares the change of directionality of the velocity vectors as a function of an increasing time delay Δt. For random diffusion, displacements are uncorrelated and the V_corr_ value is 0 for all delays. For directed motion, subsequent displacement vectors within one track will be similar, V_corr_ value will therefore be above 0. The velocity autocorrelation was measured for the collective protein behavior (i.e speckles) with the parameters specified above, as well as for single particles. For the velocity autocorrelation analysis of single molecule behavior of FtsZ, FtsA and FtsN we used TrackMate. In a first filtering step, using Track Mate’s quality threshold value, only particles exceeding quality values of 2000 were used for further analysis. Tracking was performed using the following parameters: estimated particle diameter of 0.7 μm and maximal linking distance of 0.5 μm, whereas gap-closing maximal distance was of 0.5 μm; Maximal frame gap = 2. To exclude transient, non-specific binding, only molecules present for longer than 1 frame were taken into account in our analysis. Additionally, trajectories displaying a total displacement length below 100 nm were discarded to remove non-specifically stuck particles.

### FRAP

For FRAP experiments we allowed proteins to self-organize on supported lipid membranes containing either 0.2 mol-% of tris-Ni-NTA or 1 mol-% of mono-Ni-NTA. In FRAP experiments FtsN_cyto_His, the peptide was added from bulk to steady state FtsZ pattern, as described above. FRAP experiments were performed with a frame rate of 2s/frame, and bleached area of about 15 μm × 15 μm. To obtain the half-time of fluorescence recovery, we fitted a mono-exponential function to the intensity trace using a Python-based Jython macro (Image Processing School Pilsen 2009).

To discriminate between fluorescence recovery via protein exchange or lateral diffusion a rectangular area was bleached, and fitted a 1D diffusion equation to the corresponding intensity profile: 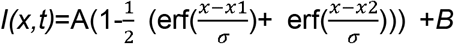, where *erf* denotes the Gaussian error function, and 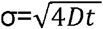. The initial height of the well was given by the intensity profile at time zero after bleaching, normalized by the background signal. In case of FtsZ, only the first 10 s of the recovery movie were fitted, while for FtsA and FtsN first 100 s were taken into consideration. The mode of recovery is revealed by a linear function σ^2^=4Dt, where an increase in the slope σ^2^ in time allows to determine the lateral diffusion coefficient. For the membrane-attached peptides, experiments required a stable peptide immobilization *via* a tris-Ni-NTA membrane linker. The comparison of the recovery half-time at the edge vs center revealed much faster recovery at edge, confirming that our immobilization strategy allows us to probe only lateral diffusion. Accordingly, σ^2^ increased with time consistent with recovery solely due lateral diffusion.

### Single molecule tracking

Single molecule tracking of FtsN_cyto_His, FtsA and FtsZ was performed with Cy5 labeled proteins in one channel and a pattern of FtsZ-Alexa488 in the other channel. For dual-color movies we acquired the two channels sequentially to minimize cross-talk.

The localization and tracking of single molecules was performed either using TrackMate (for lifetime analysis) or SLIMfast (for diffusion and confinement analysis)^11^

### FtsZ life-time analysis on single particle level

To obtain the residence time of FtsZ, unlabeled FtsA and Alexa488 FtsZ were mixed at standard concentrations (0.5 μM /1.5 μM respectively) and FtsZ-Cy5 was added to a concentration of 0.25 nM. To quantify FtsZ residence times in the presence of FtsN, 1 μM of unlabeled FtsN_cyto_His peptide was added to the experiment and the acquisition was repeated. Single molecules were tracked with TrackMate, using the same parameters as described above (“Single particle velocity autocorrelation”). Molecules only transiently present (1 frame) and longer than 100 frames due to non-specific binding were not taken into account for the analysis. The lifetime of FtsZ monomers on membrane was quantified by a monoexponential fit 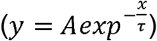 to the lifetime histograms from the reconstructed trajectories. To account on photobleaching, measured residence time was corrected by subtracting an estimated photobleaching rate of ~ 0.6±0.2 s^−1^ (ref. 12). In order to determine the photobleaching rate, FtsZ residence time was measured at four different acquisition rates (0.23 s, 0.5 s, 1 s and 2 s) at constant exposure times of 50 ms. The measured residence time was plotted against the acquisition rates and fitted with linear regression to quantify photobleaching rate.

### Diffusion analysis

Single particle localization was done in SLIMfast software using a localization module ^13^. Diffusion of membrane proteins and peptides was analyzed using SLIMfast. In this software, the tracking was performed in 2 phases: the first phase linking nearest-neighbors, while in the next phase fragments were linked into final trajectories. A Kalman filter was used to estimate the search radius within the user-defined parameters. The search radius was adapted to local density for frame-to-frame linking and final track assembly.

To quantify the diffusion coefficient a MSD-Time-Curve was calculated for each individual trajectory. Only trajectories longer than 10 frames were taken into account. The probability density distribution of diffusion coefficients calculated for FtsZ and FtsA was fitted with a Gaussian distribution. In case of FtsN a double-Gaussian fit was applied.

### Analysis of confinement time

To a steady-state pattern of FtsZ/FtsA, unlabeled FtsNcytoHis was added from bulk solution at concentration 2 μM supplemented with Cy5-labelled peptides FtsN_cyto_His or FtsN_cyto-DDEE_His at 10-20 pM. Images were acquired at 32 or 51 ms, with exposure times of 30 and 50 ms respectively.

The trajectory set produced by SLIMfast revealed heterogeneous behavior of the FtsNcytoHis peptide within a single trajectory, where fast motions were alternating with the confinement to FtsA/FtsZ filaments. Such capturing behavior was shorter for FtsN_cyto-DDEE_**His**. To analyze the contribution of confinement to the apparent diffusion coefficient, the MSD was calculated for each trajectory and fitted by a linear fit over lag 2 −10. Only trajectories of 10 frames long were taken into consideration. The distribution of diffusion coefficients was analyzed using a two-component Gaussian function to estimate weighted contribution for each population. The slow population of FtsN_cyto_**His** had more confined trajectories in comparison to FtsN_cyto-DDEE_**His**. To discard contribution of non-specifically bound molecules, only trajectories of the fast population were taken into consideration for further analysis (Supplementary Fig. 5).

The displacement distribution of the fast population for FtsN_cyto_**His** and FtsN_cyto-DDEE_**His** peptides was fitted with two-component Rayleigh distribution and weighted contribution of short and long displacements was estimated for each peptide (Fig. 3d).

The confined regions within each trajectory were identified by spatial clustering, based on DBSCAN algorithm^14^ implemented into SLIMfast software^15^. To this end a spatial cluster with a search area of 200 nm, minimal time window of 10 frames and initial window of 100 frames (at 51ms/frame acquisition rate) was built. Afterwards, the life-time of the identified clusters was fitted to a single exponential including an y-offset 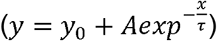 to quantify an apparent confinement time.

### Autocorrelation Analysis

To quantify the reorganization of FtsZ/FtsA filament patterns by autocorrelation we used the ImageJ plugin “Image CorrelationJ”^8^. For the analysis, we allowed the system to reach steady state and acquired time-lapse movies at a frame rate of 2 s/frame. Contrast was normalized and enhanced prior to analysis and movies aligned using the “Linear Stack Alignment with SIFT” plugin to eliminate X-Y drift. We then constructed a subset of these movies consisting of only every 10^th^ frame. This movie was then used to calculate the correlation coefficient for increasing delay Δt between two frames (“Increasing sequence”).

### FtsZ intensity measurements

To measure the intensity changes of FtsZ upon addition of peptide, no preprocessing of the movies was performed, as normalizing the intensity would lead to a loss of information. The intensity was measured with the “Stack-Z profile” function of ImageJ of the whole field of view (55 x 55 μm) of a continuous time-lapse experiment for 5 minutes before and after the peptide addition.

---

**Supplementary Figure 1.**
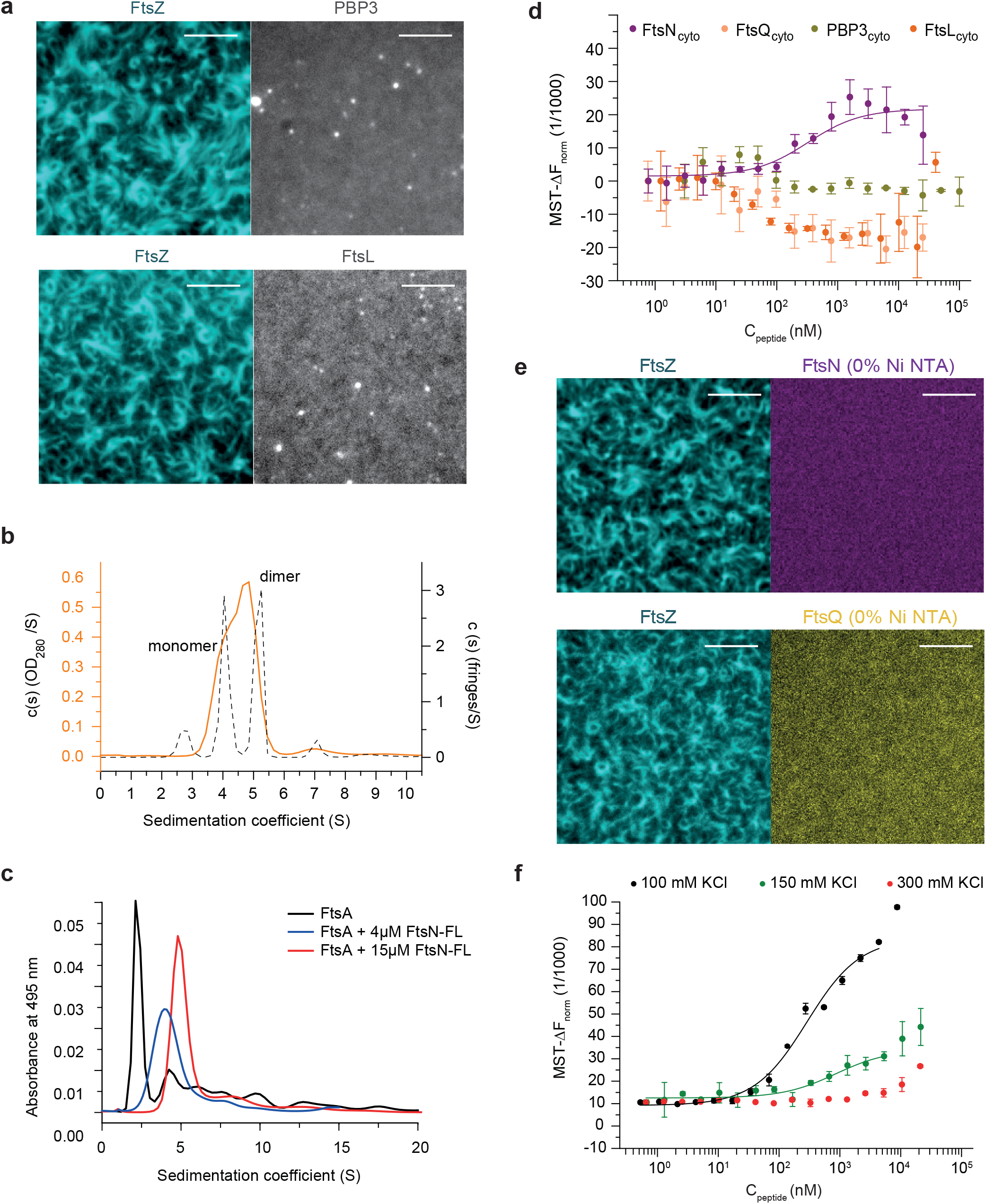

**Supplementary Figure 2.**
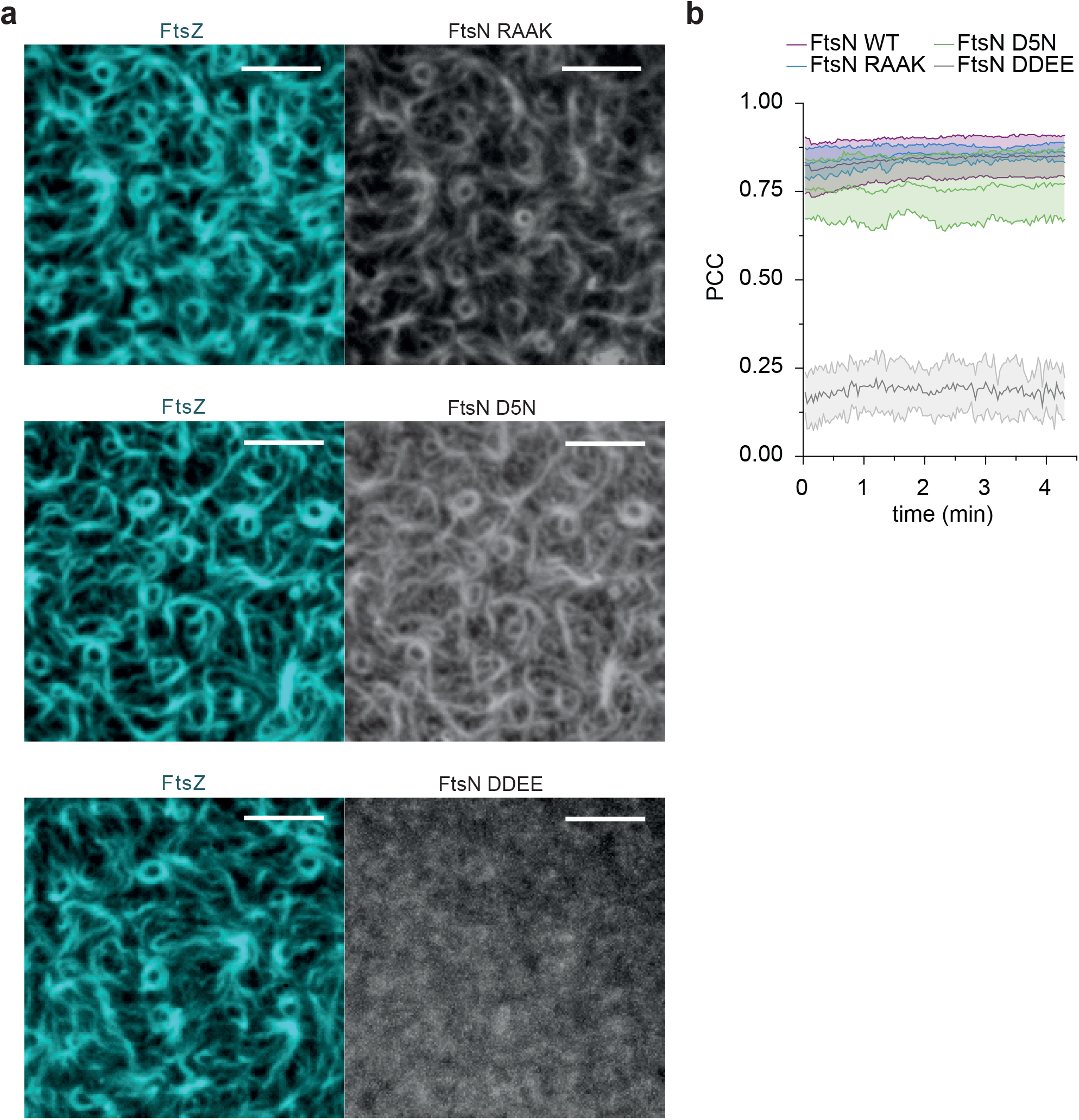

**Supplementary Figure 3.**
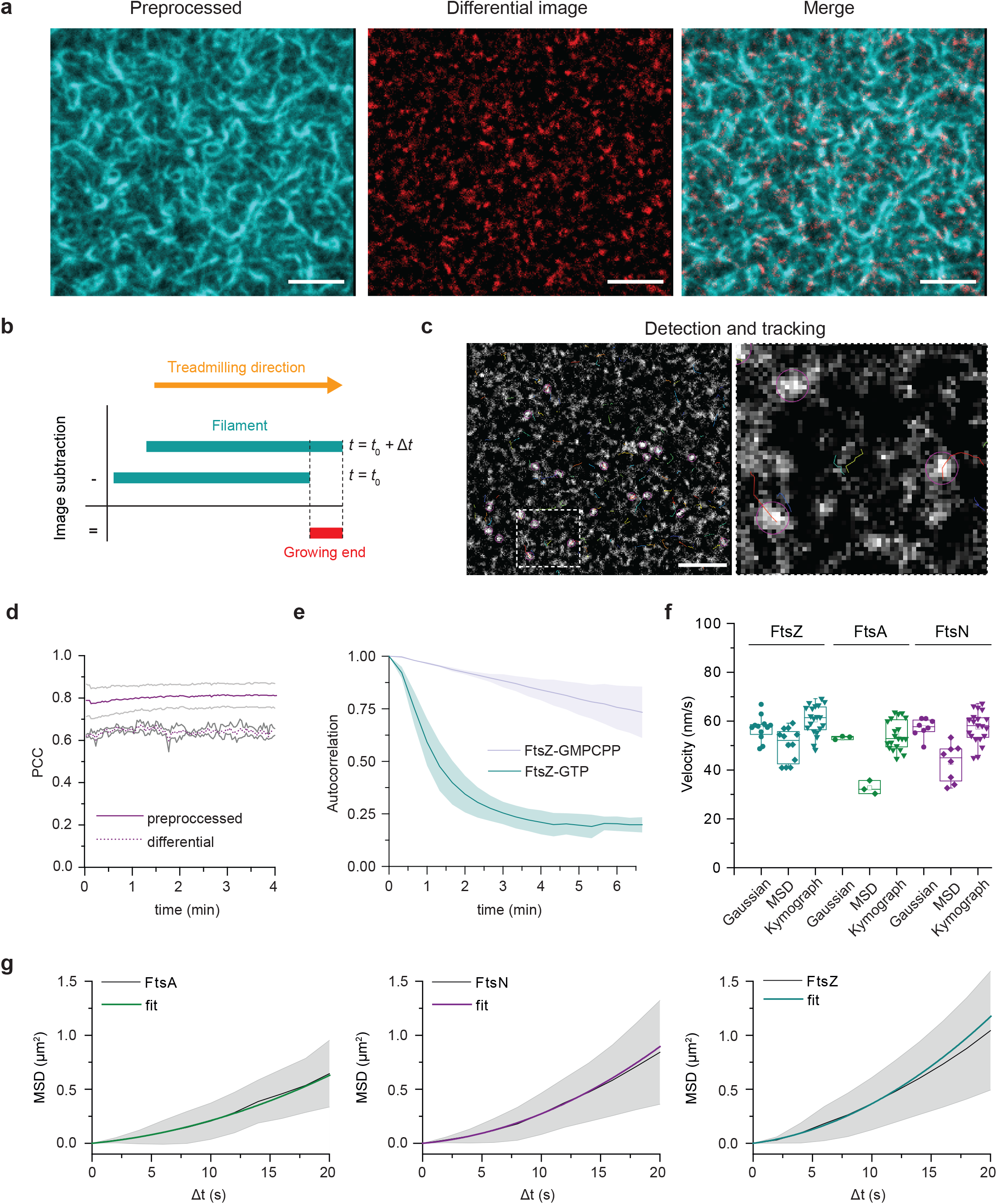

**Supplementary Figure 4.**
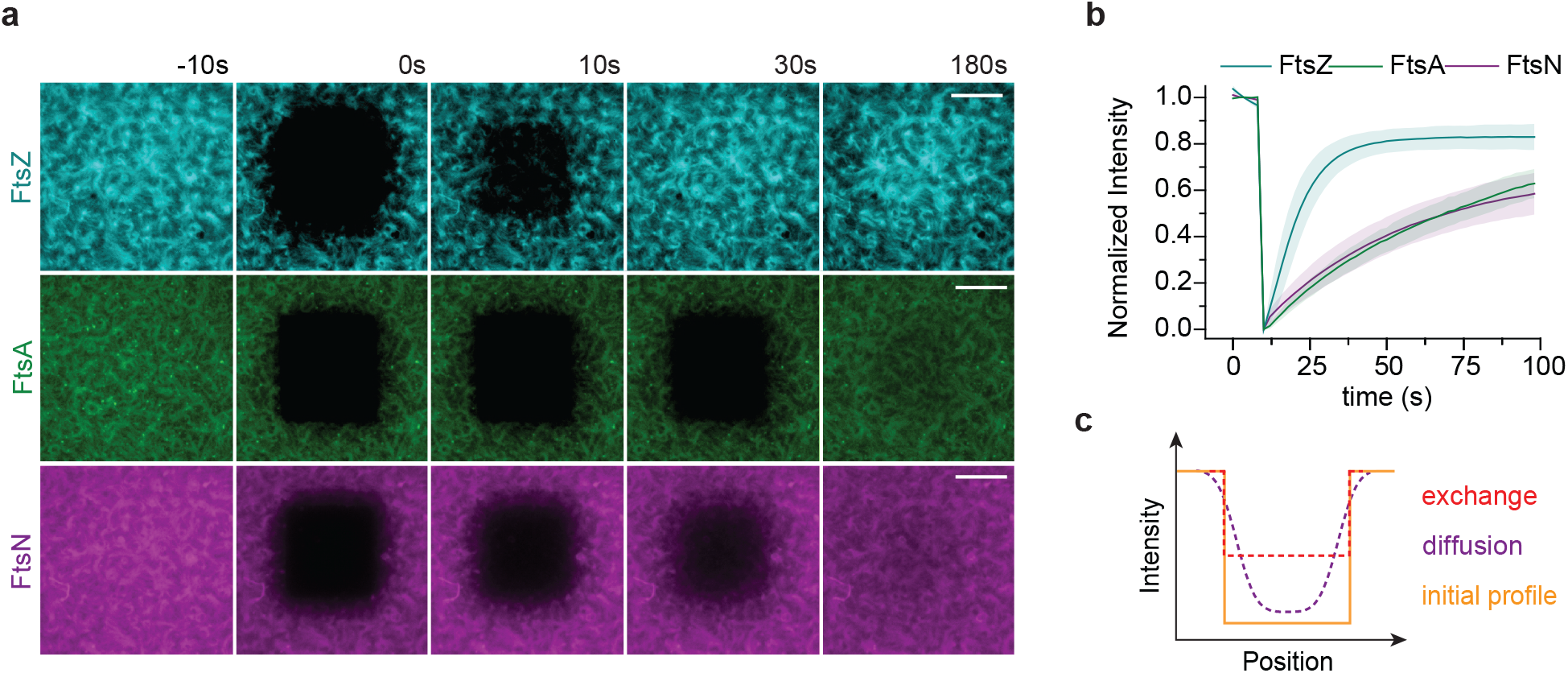

**Supplementary Figure 5.**
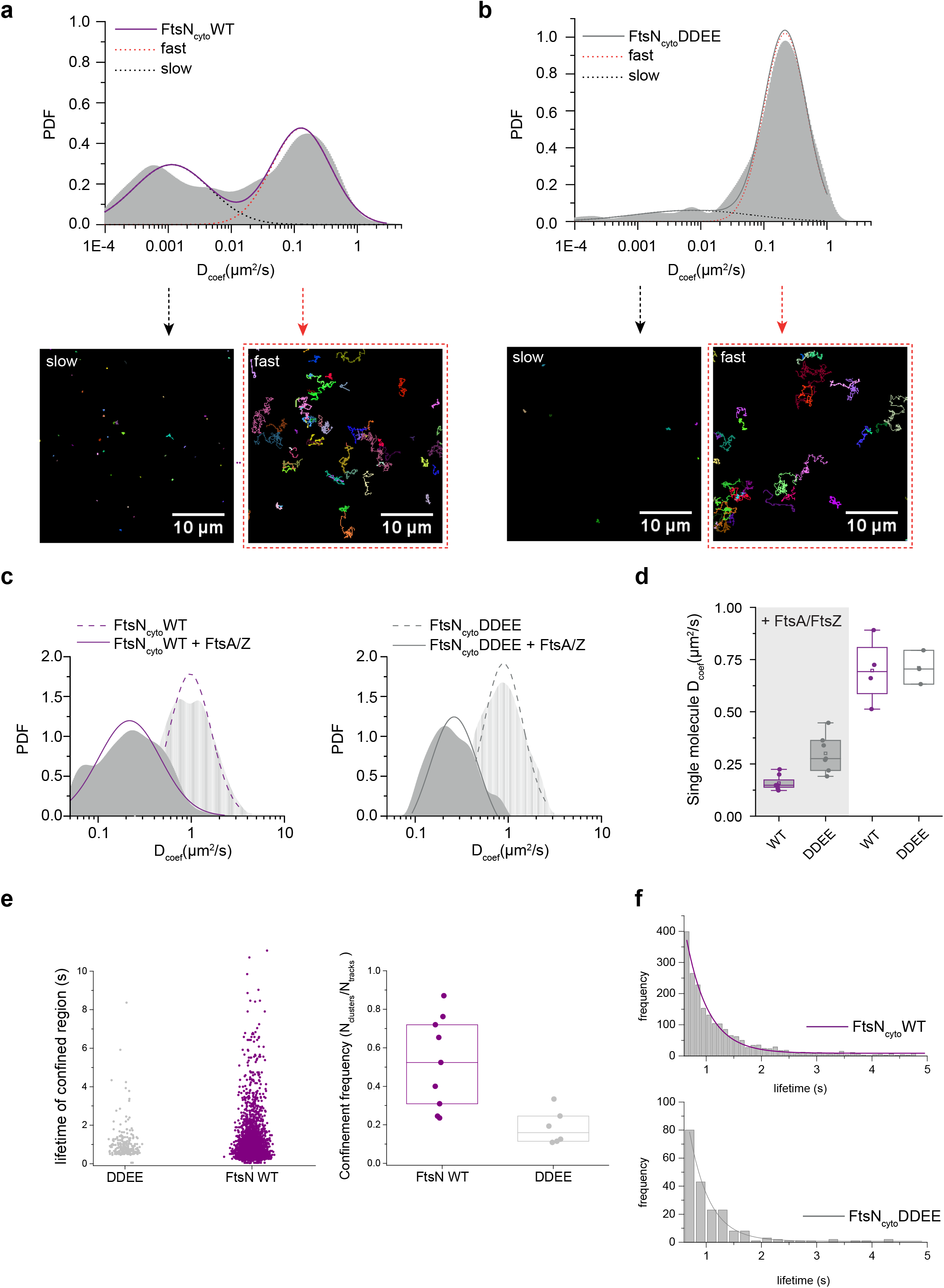

**Supplementary Figure 6.**
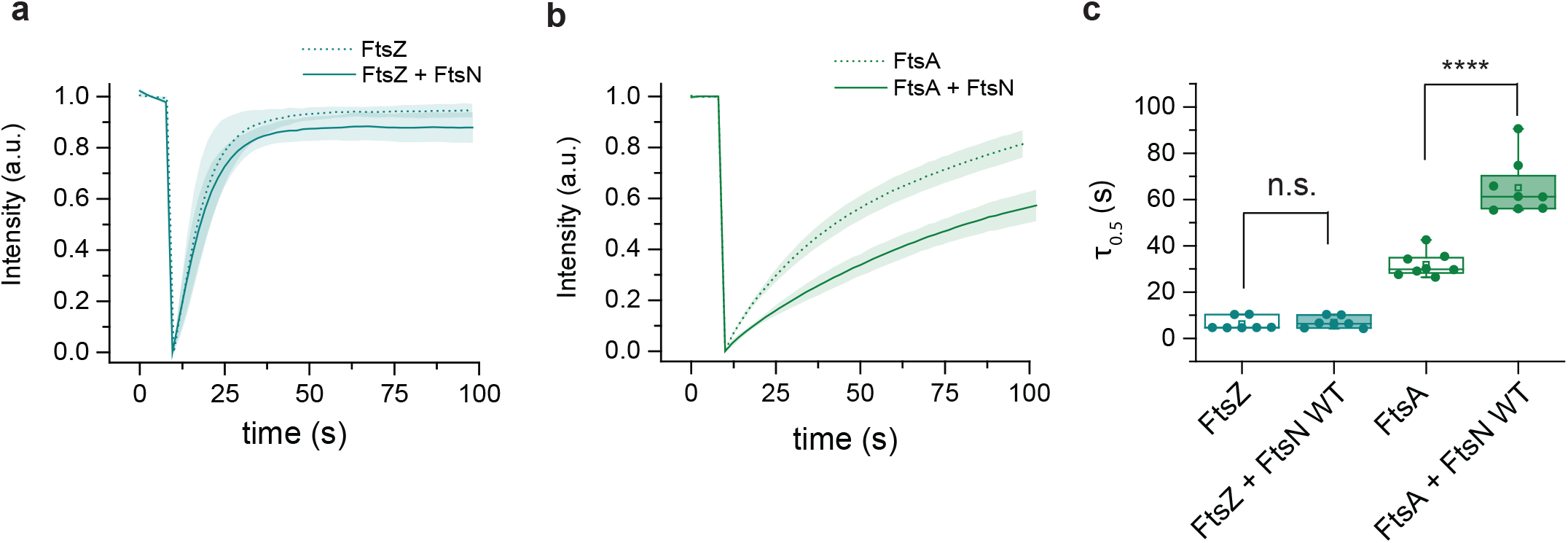

