## Supplementary material for "FtsZ assembles the bacterial cell division machinery by a diffusion-and-capture mechanism"

### Supplementary Tables

#### Supplementary Table 1: Peptide sequences

| Peptide | Sequence | MW (Da) | Purity (%) |
|---|---|---|---|
| FtsN$_{cyto}$His | CMAQR**D**YVRRSQPAPSRRKKSTSRKKQRNLPAVHHHHHH | 4691,37 | 86,51 |
| PBP3$_{cyto}$His | CMKAAAKTQKPKRQEEHANHHHHH | 2855,23 | 95,81 |
| FtsL$_{cyto}$His | CMISRVTEALSKVKGSMGSHERHALPGVIGDDLLRHHHHHH | 4587,28 | 96,90 |
| FtsQ$_{cyto}$His | CMSQAALNTRNSEEEVSSRRNNGTRHHHHHH | 3633,89 | 96,95 |
| FtsN$_{cyto-D5N}$His | CMAQR**N**YVRRSQPAPSRRKKSTSRKKQRNLPAVHHHHHH | 4723,20 | 98,41 |
| FtsN$_{cyto-RAAK}$His | CMAQRDYVRRSQPAPS**RAAK**STSRKKQRNLPAVHHHHHH | 4582,20 | 97,67 |
| FtsN$_{cyto-DDEE}$His | CMAQRDYVRRSQPAPS**DDEE**STSRKKQRNLPAVHHHHHH | 4644,72 | 95,69 |

#### Supplementary Table 2: Summary of the binding constants for different FtsN constructs to FtsA.

| FtsN Version | $K_D$ (nM) | $R^2$ |
|---|---|---|
| Full-length FtsN | 250 ± 50 | 0.988 |
| FtsN$_{cyto}$His | 324 ± 96 | 0.967 |
| FtsN$_{cyto-D5N}$His | 165 ± 30 | 0.991 |
| FtsN$_{cyto-DDEE}$His | 815 ± 81 | 0.972 |

### Supplementary Figures

**Supplementary Figure 1: Intrinsically weak interactions of $FtsN_{cyto}$His and $FtsQ_{cyto}$His peptides with FtsA in solution are enhanced by immobilization to a supported lipid membrane.**

**a.** $PBP3_{cyto}$His (top) and $FtsL_{cyto}$His (bottom) remain homogeneously distributed (n = 3) in the presence of FtsA/FtsZ filaments (cyan). Scale bars 5 µm.

**b.** Sedimentation velocity (SV) characterization of protein-detergent complexes formed by the solubilization of 16 µM full-length FtsN in 3.72 mM DDM. Sedimentation velocity c(s) distributions obtained from the analysis of the absorbance (orange line, scale $OD_{280}$ /S) and Raleigh interference (grey dashed line, scale fringes/S) signals of FtsN complexes using the program SEDFIT. Main peaks at 4.1 ± 0.1 and 5.2 ± 0.1 S detected by Raleigh interference (short dash line) are compatible with FtsN monomers and dimers.

**c.** SV analysis of Alexa(Fluor)488-FtsA in the presence of full-length FtsN at a 1:1 FtsA:FtsN molar ratio (blue) and a ~1:4 FtsA:FtsN molar ratio (red), or in the absence of FtsN (black). The main peak of FtsA at ~2 S shifts to a higher *s*-value upon addition of FtsN, indicating the formation of higher molecular weight species.

**d.** Binding curves obtained from the MST thermograms of Alexa647-FtsA titrated with $FtsN_{cyto}$His, $FtsQ_{cyto}$His, $PBP3_{cyto}$His or $FtsL_{cyto}$His. Binding was observed only for $FtsN_{cyto}$His, fitting a Hill equation gave an apparent $K_D$ of 324 ± 96 nM. The negative shift in MST curves for $FtsL_{cyto}$His and $FtsQ_{cyto}$His peptides do not represent specific binding. (See *Supplementary Table 2*).

**e.** Attachment to the membrane surface is necessary for colocalization of the cytoplasmic peptides of FtsN and FtsQ with FtsZ/FtsA co-filaments. $FtsN_{cyto}$His (magenta, A) and $FtsQ_{cyto}$His (yellow, B) added to a concentration of 1 µM do not interact with FtsA/FtsZ filaments (cyan) on a membrane without Ni-NTA. Scale bar 10 µm.

**f.** Binding curves obtained from the MST thermograms of Alexa647-FtsA titrated with full length FtsN at 100 (black symbols), 150 (green symbols) or 300 mM KCl (red symbols). The binding curves at the lower ionic strengths are biphasic with a first transition from 100 to 1000 nM FtsN and a second transition that could not be saturated. The interaction was significantly weaker in the presence of 300 mM KCl (red symbols). The Hill equation with Hill coefficient fixed to 1 was used to calculate the midpoint of the first transition for the curves at 100 mM (black line) or 150 mM KCl (green line), resulting in apparent $K_D$ values of 250 ± 51 nM at 100 mM KCl or 734 ± 23 nM at 150 mM KCl.

**Supplementary Figure 2: Ionic nature and specificity of the FtsN-FtsA interaction**

**a.** FtsN mutants (FtsN$_{cyto-RAAK}$His, FtsN$_{cyto-D5N}$His) are efficiently colocalized to FtsA/FtsZ filaments, while FtsN$_{cyto-DDEE}$His shows weak colocalization. Scale bars are 5 µm.

**b.** Colocalization of FtsN peptides with FtsZ/FtsA cofilaments is stable over time. PCC coefficients were averaged from 5 experiments, shaded areas represent standard deviations.

**Supplementary Figure 3: Differential time-lapse analysis**

**a.** Representative micrographs showing the pre-processed pattern of FtsZ (cyan) and a differential image after subtraction of a frame with a time delay of 10 s (red). Scale bars are 5 µm. Supplementary Video 5.

**b.** A schematic illustration of the image subtraction procedure.

**c.** The remaining intensity signal after differential subtraction can be detected and tracked using single particle tracking algorithms. Scale bar - 5 µm.

**d.** FtsZ/FtsN$_{cyto}$His crosscorrelation for the preprocessed images (solid magenta line, PCC = 0.84 ± 0.1 (n = 5)), and differential time lapse (dashed magenta line, PCC = 0.67 ± 0.02, (n = 4)), confirming co-migration of both proteins.

**e.** Presence of GMPCPP inhibits filament reorganization as shown by the much slower decay of the temporal autocorrelation in comparison to GTP.

**f.** Comparison of different methods for treadmilling analysis. Differential kymographs improve the accuracy of velocity determination; however, they rely on manual quantification and the sample size is smaller. Single particle tracking provides high throughput velocity quantification, where either a mean-squared displacement (MSD) or Gaussian fit to the velocity distribution can be used to quantify the treadmilling velocity. Each data point of MSD and Gaussian represents the average velocity calculated from all tracks within one independent experiment. The total number of tracks for all experiments was 8711 for FtsZ, 823 for FtsA and 5628 for FtsN$_{cyto}$His. Data points for kymographs correspond to individual ring-like FtsZ structures.

**g.** MSD analysis shows that FtsA, FtsN$_{cyto}$His and FtsZ move directionally on the ensemble level over a lag-time of 20 s.

**Supplementary Figure 4: FtsZ, FtsA and FtsN$_{cyto}$His show different diffusive behavior as revealed by FRAP analysis**

**a.** Montage shows fluorescence recovery of the patterns for FtsZ (cyan), FtsA (green) and FtsN$_{cyto}$His (magenta). Scale bars are 10 µm. Supplementary Video 7.

**b.** Recovery dynamics are different for FtsZ, FtsA and FtsN$_{cyto}$His. Pre-bleached intensity was normalized to 1 for each protein. Half-recovery time values ($T_{0.5}$ ) are 7.53 ± 3.51 s (n = 10) for FtsZ, 64 ± 15 s (n = 3) for FtsA and 27.61 ± 10 s (n = 10) for FtsN$_{cyto}$His.

**c.** Schematic illustration of the intensity profiles across a rectangular photobleached region (solid orange line) during recovery due to lateral 2D diffusion (magenta dashed line) or exchange with labeled proteins in solution (red dashed line).

**Supplementary Figure 5: Single-molecule analysis of FtsN$_{cyto}$His diffusion-and-capture by FtsA/FtsZ filaments**

**a,-b.** Probability density distributions of the diffusion coefficients quantified for each single-molecule trajectory of FtsN$_{cyto}$His (a) and FtsN$_{cyto\text{-}DDEE}$His (b). Single-molecule tracking of diluted Cy5-labelled peptides was performed in the presence of FtsA/FtsZ filaments and standard concentration of unlabeled peptides. Gaussian fits of the distributions revealed the presence of two populations: a slow population with $D_{coef}$ = 0.0011 ± 0.0003 μm$^2$/s and a fast population with $D_{coef}$ = 0.15 ± 0.3 μm$^2$/s for FtsN$_{cyto}$His (left), and $D_{coef}$ = 0.3 ± 0.1 μm$^2$/s for FtsN$_{cyto\text{-}DDEE}$His (right). The slow populations for both peptides contain both immobile particles and molecules with confined diffusion (dashed arrow points to corresponding trajectory maps for slow and fast populations).

**c.** The mean diffusion coefficient of single molecules of FtsN without FtsA/FtsZ filaments was comparable between FtsN$_{cyto}$His ($D_{coef}$ = 0.69 ± 0.16 μm$^2$/s, n = 5, left) and FtsN$_{cyto\text{-}DDEE}$His ($D_{coef}$ = 0.71 ± 0.10 μm$^2$/s, n = 3, right), (light-grey histogram, dashed lines). Addition of FtsA/FtsZ proteins slowed down diffusion of both peptides: FtsN$_{cyto}$His ($D_{coef}$ = 0.16 ± 0.04 μm$^2$/s, n = 8) and FtsN$_{cyto\text{-}DDEE}$His ($D_{coef}$ = 0.30 ± 0.08 μm$^2$/s, n = 7), (dark-grey histogram, solid lines).

**d.** The mean squared diffusion coefficients are comparable between FtsN$_{cyto}$His and FtsN$_{cyto\text{-}DDEE}$His without FtsA and FtsZ proteins present (white boxes). FtsA/FtsZ filaments on the membrane slow down diffusion of both peptides (grey boxes), but to a different extent: diffusion of FtsN$_{cyto}$His is 4.4-fold slower, while diffusion of FtsN$_{cyto\text{-}DDEE}$His is slowed down by 2.4-fold.

e. The fast-diffusing particle populations (Fig. 5a-c) for the wild type FtsN$_{cyto}$His and DDEE mutant were subjected to further clustering analysis to identify the regions of the trajectories when confined motion happens using a clustering algorithm (as described in the methods). Left: The trajectories of wild type FtsN$_{cyto}$His particles contained more confined motion events in comparison to the DDEE mutant. Each dot in the scatter plot represents a single region of confined motion detected by the algorithm in all trajectories from 3 independent experiments. Right: The number of confined regions was normalized by the total amount of trajectories to visualize the difference in the frequency of confinement periods between the two peptides.

f. The lifetime of the confinement was determined from a monoexponential fit to lifetime histograms. FtsN$_{cyto}$His shows slightly longer confinement (0.41±0.02s, $N_{clusters}$=1827, $R^2$= 0.985) in comparison to the DDEE mutant (0.36±0.02s, $N_{clusters}$=283, $R^2$= 0.987).

**Supplementary Figure 6: Effect of FtsN$_{cyto}$His WT addition on the turnover rate of FtsZ and FtsA.**

**a.** Addition of FtsN$_{cyto}$His from bulk solution to FtsA/FtsZ filaments does not affect FtsZ turnover on the membrane. The FRAP recovery curve for FtsZ without FtsN$_{cyto}$His (dashed line) is comparable to the curve obtained in the presence of FtsN$_{cyto}$His (solid line).

**b.** In contrast, addition of FtsN$_{cyto}$His slows down FtsA recovery time..

**c.** Recovery half-time after photobleaching of FtsA is significant decreased in the presence of FtsN$_{cyto}$His (39 ± 14 s vs. 64 ± 15 s, n = 3), while FtsZ recovery time is not affected (6.22 ± 2.77 s vs. 6.84 ± 2.44 s, n=3). Recovery half-times for FtsZ (cyan symbols) or FtsA (green symbols) were determined in the presence (filled boxes) or absence (empty boxes) of FtsN$_{cyto}$His. Each dot represents a different position on the sample acquired in 3 independent experiments.

**Supplementary Video Legends**

**Supplementary Video 1: Membrane-bound FtsN$_{cyto}$His is sorted by co-filaments of FtsA and FtsZ**

FtsN$_{cyto}$His is homogeneously distributed on a membrane when no other proteins are present (0-30s, magenta, right). After addition of FtsA and FtsZ to the reaction chamber (noisy signal in both channels), FtsZ filaments assemble on the membrane (cyan, left). Within 5 minutes after the addition, FtsN$_{cyto}$His is sorted into a pattern resembling the FtsZ filaments.

The experiment was performed on 1 mol% Mono-Ni-NTA membranes with 1.5 μM Cy5-FtsZ, 0.5 μM FtsA and 1 μM CF488-FtsN$_{cyto}$His. The movie was acquired at 2 s/frame and corresponds to the micrographs on Fig. 1b.

**Supplementary Video 2: Membrane-bound FtsQ$_{cyto}$His is sorted by co-filaments of FtsA and FtsZ**

FtsQ$_{cyto}$His is homogeneously distributed on a membrane prior to addition of FtsA and FtsZ (0-1min, yellow, middle). After addition of FtsA and FtsZ to the reaction, FtsZ filaments assemble on the membrane (cyan, left). Within several minutes, membrane-bound FtsQ$_{cyto}$His is organized into a pattern resembling FtsZ filaments.

The experiment was performed on 0.2 mol% Tris-Ni-NTA membranes with 1.5 μM Alexa488-FtsZ, 0.5 μM FtsA and 1 μM Cy5-FtsQ$_{cyto}$His. Acquisition rate was 2s/frame. The movie corresponds to the micrographs on Fig. 1c.

**Supplementary Video 3: FtsN$_{cyto}$His and FtsQ$_{cyto}$His are simultaneously sorted by co-filaments of FtsA and FtsZ**

FtsN$_{cyto}$His and FtsQ$_{cyto}$His are simultaneously bound and homogeneously distributed on the membrane before addition of FtsA-FtsZ (0-1min, cyan and yellow respectively). After the addition of FtsA and FtsZ, FtsN$_{cyto}$His and to FtsQ$_{cyto}$His peptides are co-localized into a filament-like patterns.

The experiment was performed on 0.5 mol% Tris-Ni-NTA membranes with 0.15 µM Alexa488-FtsN$_{cyto}$His, 0.75 µM Cy5-FtsQ$_{cyto}$His, 0.5 µM FtsA and 1.5 µM FtsZ. The movie was acquired at 2 s/frame and corresponds to Fig. 1d.

**Supplementary Video 4: Colocalization of FtsN to FtsZ is mediated via FtsN$_{cyto}$ interaction with FtsA**

FtsN$_{cyto}$His shows similar colocalization to FtsZ ( left) and FtsA (right). Top: CF488-FtsN$_{cyto}$His (magenta, middle) was added to FtsA/Cy5-FtsZ co-filaments at the steady state (cyan, left). A merge of both channels displays the high efficiency of colocalization between CF488-FtsN$_{cyto}$His and treadmilling filaments of FtsZ (bottom left).

Bottom: CF488-FtsN$_{cyto}$His (magenta, -top right) was added to TMR-FtsA/FtsZ co-filaments at the steady state (green, top left). A merge of both channels displays the high amount of colocalization between CF488-FtsN$_{cyto}$His and the co-treadmilling filaments of FtsA (bottom right). The experiments were acquired at 2 s/frame and correspond to data on Fig. 1e-g.

**Supplementary Video 5: Differential imaging of treadmilling dynamics**

A merge of FtsZ and FtsN$_{cyto}$His pattern is presented on the top left, while the bottom left represents a merge of the differential image of FtsZ and FtsN$_{cyto}$His. The differential movie shows that FtsN$_{cyto}$His follows the directional motion of treadmilling FtsZ closely. The right side presents an overlay of FtsN$_{cyto}$His (cyan) on top of the FtsZ pattern (magenta), displaying that speckles corresponding to growing filament bundles move along the filaments.

The experiment was performed on a 1 mol% mono-Ni-NTA membranes with 0.5 µM FtsA, 1.5 µM Cy5-FtsZ and 1 µM CF488-FtsN$_{cyto}$His and acquired at 2 s/frame. The differential movie was created as described in the methods section. The movie corresponds to Supplementary Fig. 3a.

**Supplementary Video 6: Colocalization of FtsN$_{cyto}$His to FtsA/FtsZ filaments and the influence of treadmilling dynamics**

CF488-FtsN$_{cyto}$His shows stronger colocalization to FtsZ in the presence of GTP (top row) than in the presence of GMPCPP (bottom row).

Top: CF488-FtsN$_{cyto}$His (magenta, right) was added to the reaction chamber with FtsA and FtsZ co-filaments at steady state (cyan, left).
Bottom: CF488-FtsN$_{cyto}$His (magenta, right) was added to a reaction chamber containing 0.5 µM FtsA and 1.5 µM Cy5-FtsZ (cyan, left).
In both experiments, 0.5 µM FtsA, 1.5 µM Cy5-FtsZ and 1 µM CF488-FtsN$_{cyto}$His, as well as 4 mM GTP and 2 mM GMPCPP, respectively were used. For GTP experiments, the acquisition rate was kept at 2s/frame, while the rate was decreased in the presence of GMPCPP to 5 s/frame to limit photobleaching. The raw files were processed to show every 5$^{th}$ (GTP) or every 2$^{nd}$ (GMPCPP) frame, resulting in a 10 s/frame movie. The movies correspond to Fig. 2c.

**Supplementary Video 7: FRAP**

FtsZ, FtsA and FtsN$_{cyto}$His dynamics arise from different biochemical processes, as shown by FRAP experiments on supported lipid bilayers. The difference in the recovery times of the patterns is evident, with FtsZ showing the fastest recovery, mainly by exchange with FtsZ in the buffer and no influence of diffusion on the membrane, while fluorescence recovery of FtsA and FtsN$_{cyto}$His is slower and dominated by lateral diffusion.
Each experiment was performed in the presence of 1.5 µM FtsZ, 0.5 µM FtsA and 1 µM FtsN$_{cyto}$His labelled with Cy5, TMR and CF488CF, respectively. All movies were acquired at 2 s/frame and correspond to Fig. 2g.

**Supplementary Video 8: Comparison of single particle dynamics**

Top: Single molecule dynamics of Cy5-FtsZ, Cy5-FtsA and Cy5-FtsN$_{cyto}$His. Bottom: Single molecule dynamics merged with the corresponding Alexa488-FtsZ pattern.
Single molecules of FtsZ display no diffusive behavior at all, monomers within the filaments are constantly exchanged by binding/unbinding, as expected for treadmilling filaments. Single molecules of FtsA display a mixed behavior of lateral diffusion and binding/unbinding. Single molecules of FtsN move by fast lateral diffusion on the plane of the membrane, showing transient confinement to FtsA/FtsZ co-filaments.
Each experiment was performed on 0.2 mol% Tris-Ni-NTA membranes with 1.5 µM Alexa488-FtsZ, 0.5 µM FtsA and 1 µM FtsN$_{cyto}$His present. For visualization of the single particles, small amounts of Cy5-labeled FtsZ, FtsA or FtsN$_{cyto}$His was added (100 pM, 75 pM and 75 pM respectively). The movies were recorded at 0.228 s/frame and correspond to Fig. 3a.

**Supplementary Video 9: Comparison of single molecule trajectories of FtsN WT and FtsN DDEE**

Representative trajectories showing the differences in the diffusive motion of FtsN$_{cyto\text{-}DDEE}$His (gray track, left) and FtsN$_{cyto}$His (magenta, right). FtsN$_{cyto\text{-}DDEE}$His covers a large area and does not remain at isolated spots. $FtsN_{cyto}His$ shows similar diffusive motion in the beginning until it is captured by FtsZ/FtsA co-filaments where it shows confined behavior. The red links represent gaps in trajectory linking.
